# Spatial chromosome folding and active transcription drive DNA fragility and formation of oncogenic *MLL* translocations

**DOI:** 10.1101/485763

**Authors:** Henrike Johanna Gothe, Britta Annika Maria Bouwman, Eduardo Gade Gusmao, Rossana Piccinno, Sergi Sayols, Oliver Drechsel, Giuseppe Petrosino, Vera Minneker, Natasa Josipovic, Athanasia Mizi, Christian Friberg Nielsen, Eva-Maria Wagner, Shunichi Takeda, Hiroyuki Sasanuma, Damien Francis Hudson, Thomas Kindler, Laura Baranello, Argyris Papantonis, Nicola Crosetto, Vassilis Roukos

**Author notes:** These authors contributed equally (Britta Annika Maria Bouwman, Eduardo Gade Gusmao, Argyris Papantonis, Nicola Crosetto). Correspondance: Vassilis Roukos.

## Abstract

How spatial chromosome organization influences genome integrity is still poorly understood. Here we show that DNA double-strand breaks (DSBs) mediated by topoisomerase 2 (TOP2) activities, are enriched at chromatin loop anchors with high transcriptional activity. Recurrent DSBs occur at CTCF/cohesin bound sites at the bases of chromatin loops and their frequency positively correlates with transcriptional output and directionality. The physiological relevance of this preferential positioning is indicated by the finding that genes recurrently translocating to drive leukemias, are highly transcribed and are enriched at loop anchors. These genes accumulate DSBs at recurrent hot spots that give rise to chromosomal fusions relying on the activity of both TOP2 isoforms and on transcriptional elongation. We propose that transcription and 3D chromosome folding jointly pose a threat to genomic stability, and are key contributors to the occurrence of genome rearrangements that drive cancer.

## Introduction

Cellular processes, such as DNA replication, transcription and DNA condensation generate torsional stress that can be resolved by topoisomerases. Type I topoisomerases, such as TOP1, dissipate torsional stress by transiently nicking and religating one DNA strand, while type II topoisomerases (TOP2), regulate DNA topology by relaxing, unknotting and decatenating DNA by transiently breaking and rejoining both DNA strands of the double helix (Ashour et al., 2015; Pommier et al., 2016). The two TOP2 isoforms, TOP2A and TOP2B, share 70% sequence identity, but have distinct expression patterns. TOP2A is essential for viability, it is expressed predominantly in cycling cells during the S and G2/M phases of the cell cycle and plays key roles in replication, chromosome segregation and transcription (Pendleton et al., 2014). TOP2B participates mainly in transcription (Kouzine et al., 2013; Naughton et al., 2013), it is ubiquitously expressed in both cycling and postmitotic cells, and although it is dispensable for cell proliferation, it is required for neural development in mice (Ju et al., 2006; Yang et al., 2000).

How changes in the local DNA topology interface with the 3D chromosome organization is poorly understood. Hierarchical loop structures, ranging from promoter-enhancer interactions to larger topological domains, shape chromosome structure (Pombo and Dillon, 2015; Rao et al., 2014). On a local scale, chromatin is folded into loops (Rao et al., 2014) and topological associated domains (TADs) (Dixon et al., 2012; Nora et al., 2012). In mammals, loop anchors frequently contain CTCF motifs oriented in a convergent manner bound by CTCF and cohesin (de Wit et al., 2015; Rao et al., 2014), and TOP2B which physically interacts with CTCF and cohesin (Uuskula-Reimand et al., 2016), has been found enriched at these regions (Canela et al., 2017; Madabhushi et al., 2015; Uuskula-Reimand et al., 2016). It has been hypothesized therefore, that TOP2 may acts at chromatin loop anchor sites to dissipate torsional stress arising during transcription (Uuskula-Reimand et al., 2016) and topological constraints during loop extrusion dynamics (Canela et al., 2017).

Although physiologically important, TOP2 functions are also inherently risky for a cell, as they involve the formation of a transient DSB to allow the passage of duplex DNA through the break. During this controlled process, the two-TOP2 subunits are covalently linked through their active sites to each 5’-terminus of a DSB via a phosphodiester bond (Pommier et al., 2016). These key intermediates of TOP2 activity, called TOP2 cleavage complexes (TOP2ccs) are normally short-lived, as topoisomerases quickly ligate the DNA ends upon the passage of the duplex DNA. In the presence of nearby DNA lesions or upon treatment with topoisomerase poisons, however, these cleavage complexes can be stabilized on DNA and may contribute to genomic instability by acting as road blocks to DNA-tracking systems that attempt to traverse them (Ashour et al., 2015).

The mechanism by which TOP2ccs are converted to DSBs underlies the action of widely used class of anticancer agents that trap TOP2 at the intermediate cleavage complex step (Nitiss, 2009). The TOP2 poison etoposide (ETO), is among the most effective and widely used anticancer agents in the clinic, but treatment with etoposide is associated with the occurrence of therapy-related acute myeloid leukemias (t-AMLs) (Allan and Travis, 2005; Wright and Vaughan, 2014). Approximately one third of t-AML cases are associated with recurrent chromosome translocations between the mixed lineage leukemia gene (*MLL*) and various potential translocation partners, and amongst them, *AF9, AF4, AF6,* and *ENL,* account for approximately 80% of the cases. Sequencing of *MLL* fusions from patients with t-AML has shown that *MLL* breakpoints occur in breakpoint cluster regions (BCRs) at restricted genomic positions near exon 12 of the *MLL* gene, while BCRs within potential translocation partners are much broader (Zhang and Rowley, 2006). The prevailing model of ETO-induced breakage at *MLL* involves misrepair of TOP2-mediated DNA damage (Lovett et al., 2001; Povirk, 2006), but how poisoned TOP2ccs are converted to breaks at BCRs in *MLL* and translocation partner genes is unclear. It has been suggested that ETO-induced *MLL* breakage may result from direct TOP2B-mediated cleavage at *MLL* BCRs, and that active transcription is required to keep the involved partners in proximity, providing a time window for illegitimate joining and translocations (Cowell et al., 2012). In support, inhibition of transcription in immortalized retinal epithelial cells led to a decrease in *MLL* locus breakage, but a possible effect on the formation of *MLL* fusions was not assessed (Gomez-Herreros et al., 2017). Profiling of ETO-induced DSBs in mouse primary B-lymphocytes suggested that TOP2-dependent DSBs within *MLL* were largely transcription-independent, arguing against a role of transcription in ETO-induced DNA damage within *MLL* and its potential translocation partners (Canela et al., 2017). Instead, it was proposed that chromatin entanglements at topological domain borders generated during loop dynamics, require transcription-independent functions of TOP2B, promoting chromosome breakage within the *MLL* gene and the potential translocation partners (Canela et al., 2017).

Here we deployed a comprehensive approach of combining genomic and high-throughput imaging methodologies to show that transcription and chromosome folding are major contributors to TOP2-induced genomic instability and to the formation of oncogenic *MLL* fusions. We show that TOP2-induced DSBs are enriched at highly active genes located at loop anchors marked by CTCF/cohesin sites, and that DSB frequency positively correlates with transcriptional output and directionality. Likewise, transcription-dependent DSBs within translocation BCRs of highly transcribed *MLL* and fusion partner genes are localized within chromatin loop boundaries and form *MLL* fusions in a transcription- and cell cycle-dependent manner. Factors involved in removal of trapped TOP2 on DNA and the non-homologous end-joining (NHEJ) repair pathway, act as suppressors of *MLL* translocations by preventing TOP2-induced DSBs or by facilitating intrachromosomal repair within the *MLL* gene. We propose that abortive TOP2 activity, in an attempt to release transcription-dependent torsional stress at loop boundaries, generates breaks, driving the formation of oncogenic translocations.

## Results

### Genome-wide profiling of etoposide-induced DSBs in human hematopoietic cells

To determine whether *MLL* fusions are associated with DSBs at specific translocation hotspots found in t-AML patients upon ETO treatment, we profiled the ETO-induced DSBs at nucleotide resolution across the genome using suspension-cell BLISS (sBLISS), an adaptation of the BLISS methodology for mapping DSBs genome-wide (Yan et al., 2017). Since *MLL* translocations can induce acute leukemias from hematopoietic progenitor subsets (Krivtsov and Armstrong, 2007), we examined the effect of physiologically relevant doses of ETO (20 to 30μM for 4-6h) (Goodman et al., 2001), in relevant human hematopoietic erythroleukemia K562 and lymphoblastoid TK6 cells. sBLISS identified numerous ETO-induced breakage hot spots across the genome in both cell lines (K562: 10,682 hotspots; TK6 cells: 44,900 hotspots) (Figures 1A and S1A). In line with established TOP2 functions during transcription (Ju et al., 2006; Lyu et al., 2006), ETO-induced hot spots were found enriched at promoter regions (55.22%) and enhancers (15.6%), but were also distributed across gene bodies (12.34%) and intergenic regions (16.84%) (K562 cells, Figures 1B and 1C). Genome-wide correlation analysis showed that ETO-induced DSBs were highly enriched at sites of open chromatin marked by DNase I hypersensitivity (Spearman’s correlation=0.67, *p*<10^−5^) and highly correlated with RNA polymerase II occupancy (Spearman’s correlation=0.66, *p*<10^−5^) and its serine 2 (Ser2) and serine 5 (Ser5) active forms (pol2 Ser2, Spearman’s correlation= 0.57 [*p*<10^−4^] and pol2 Ser5, Spearman’s correlation=0.64 [*p*<10^−5^]) (Figures 1C,D and S1B,C). In agreement with recent observations (Canela et al., 2017; Uuskula-Reimand et al., 2016) we found that ≈40% of ETO-induced hotspots coincided with binding sites of CTCF and cohesin subunits (CTCF, SMC3, Rad21; *p*<10^−6^, Mann–Whitney–Wilcoxon test) (Figures 1C,D and S1B-D). Consistent with the observed preference for DNase I hypersensitive regions, we also found that ETO-induced DSBs are enriched at nucleosome-free regions (*p*<10^−4^, Mann–Whitney–Wilcoxon test), as revealed by MNase-seq experiments (Figures 1C,D and S1B).

**Figure 1.**
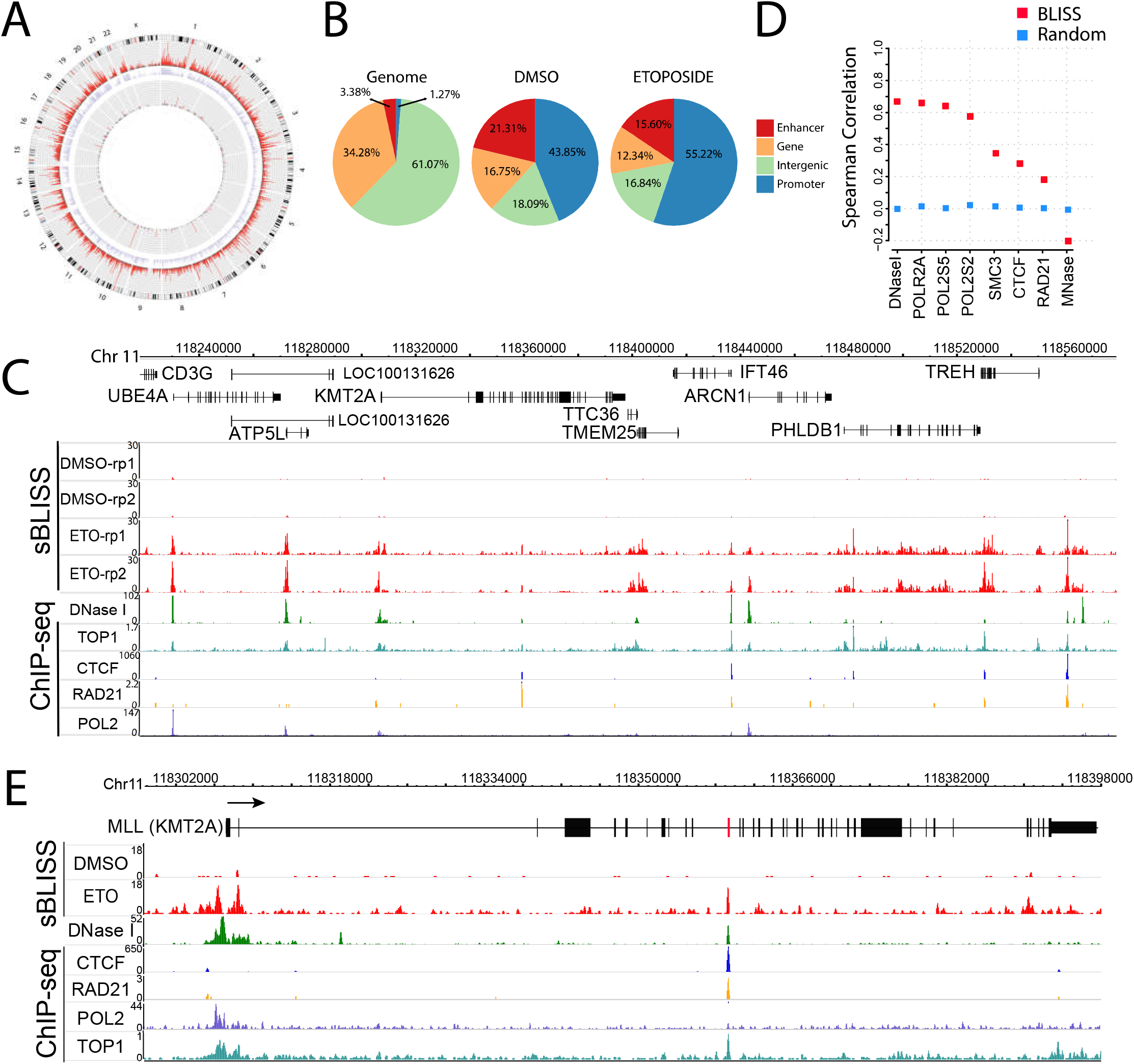
Genome-wide mapping of ETO-induced DSBs by sBLISS. (A), Circos plot depicting DSB hotspots detected by sBLISS in DMSO-(inner two circles) or ETO-treated (30μM, 4h) TK6 cells (two outer circles). Grey tracks mark hot spots with adjusted p<0.05, red tracks with adjusted p<1e-10. Coverage density of hotspots along chromosomes are marked light purple. (B) Distribution of spontaneous (DMSO) and ETO-induced (20μM, 6h) DSB hotspots in K562 cells across the genome. The distribution of the assessed features across the genome (Genome) is also shown for comparison. (C) sBLISS DSB profiles in DMSO or ETO-treated (30μM, 4h) TK6 cells shown in replicates (rp1, rp2). DNase I hypersensitivity data and occupancy of CTCF, Rad21 and Pol2 measured by ChIP-seq were obtained from ENCODE (Davis et al., 2018) and TOP1 ChIP seq from (Baranello et al., 2016). (D) Correlation analysis (Spearman’s correlation coefficient, BLISS, marked red) between ETO-induced DSBs (20μM, 6h) and open chromatin genome sites (probed by DNaseI-seq), nucleosome occupancy (probed by MNase-seq) or occupancy of Pol2, Pol2Ser2, Pol2Ser5, CTCF, Rad21, SMC3 in K562 cells (probed by ChIP-seq, all data derived from ENCODE); computed correlation of one hundred randomized tests with DSBs within the same regions of these features is also shown (random; blue). (E) sBLISS DSB-profiles along the *MLL (KMT2A)* in DMSO and ETO-treated (30μM, 4h) TK6 cells. Exons are shown as black squares, the t-AML translocation hot spot in *MLL* is shown red. ChIP-seq data as in (C), arrow represents the direction of transcription.

To check whether the observed ETO-induced DNA breakage is associated with the known translocation hotspots found in t-AML patients, we probed for enrichment of ETO-induced DSBs within the BCR of *MLL* and its potential fusion partners. ETO treatment in both TK6 and K562 human hematopoietic cell lines (Figures 1E and S1F) led to the localized formation of DSBs in the exon 12 BCR of the *MLL* gene. This ETO-induced *MLL* DSB-hotspot coincides with DNase I hypersensitivity, CTCF, cohesin and TOP2B binding and activity sites (Canela et al., 2017; Cowell et al., 2012), suggesting that TOP2ccs, stabilized by treatment with ETO at this site, may be converted to clean DSBs that fuel the formation of *MLL* fusions. ETO treatment led to an enrichment of DSBs within translocation hotspot areas of the most common *MLL* translocation partners (Figure S1G), but exhibited broader breakage patterns, suggesting that selection during oncogenesis determines the translocated areas of the potential partners involved in *MLL* fusions.

We also noticed that several of the genomic sites that exhibited high levels of ETO-induced DSBs also showed spontaneous breakage of endogenous sites in the absence of the drug (DMSO control, Figures 1A and S1A). While endogenous DSB hotspots were significantly fewer than the ETO-induced ones and varied between the different cell lines (K562, DMSO: 3,225 ETO: 10,682; TK6, DMSO: 329, ETO: 44,900, Figures 1A and S1A), 67% of the spontaneous DSB hot spots detected by sBLISS were also found in ETO-treated samples (K562 cells, Figure S1E) and exhibited a similar distribution across the genome, being mainly localized in gene promoters (Figure 1B, DMSO control). The similarity between localization patterns of endogenous and ETO-induced DSB hotspots suggests that a substantial fraction of spontaneous breaks in a cell might arise from abortive topoisomerase activity.

### *MLL* fusions form rarely in hematopoietic cell lines and progenitor cells

To estimate the frequency of ETO-induced *MLL* fusions, we developed C-Fusion 3D, a high-throughput imaging-based methodology to probe chromosome breakage and rare translocations in individual cells with high sensitivity. Built upon the rationale of high-throughput break-apart fluorescence *in situ* hybridization (FISH) (Burman et al., 2015a; Roukos et al., 2013), C-Fusion 3D detects individual cells with chromosome breakage and/or fusions by measuring the extent of spatial separation or colocalization of individual chromosome breaks in 3D (Figure 2A). We found that C-Fusion 3D can detect individual cells with chromosome translocations at frequencies down to 10^−4^ (data not shown).

**Figure 2.**
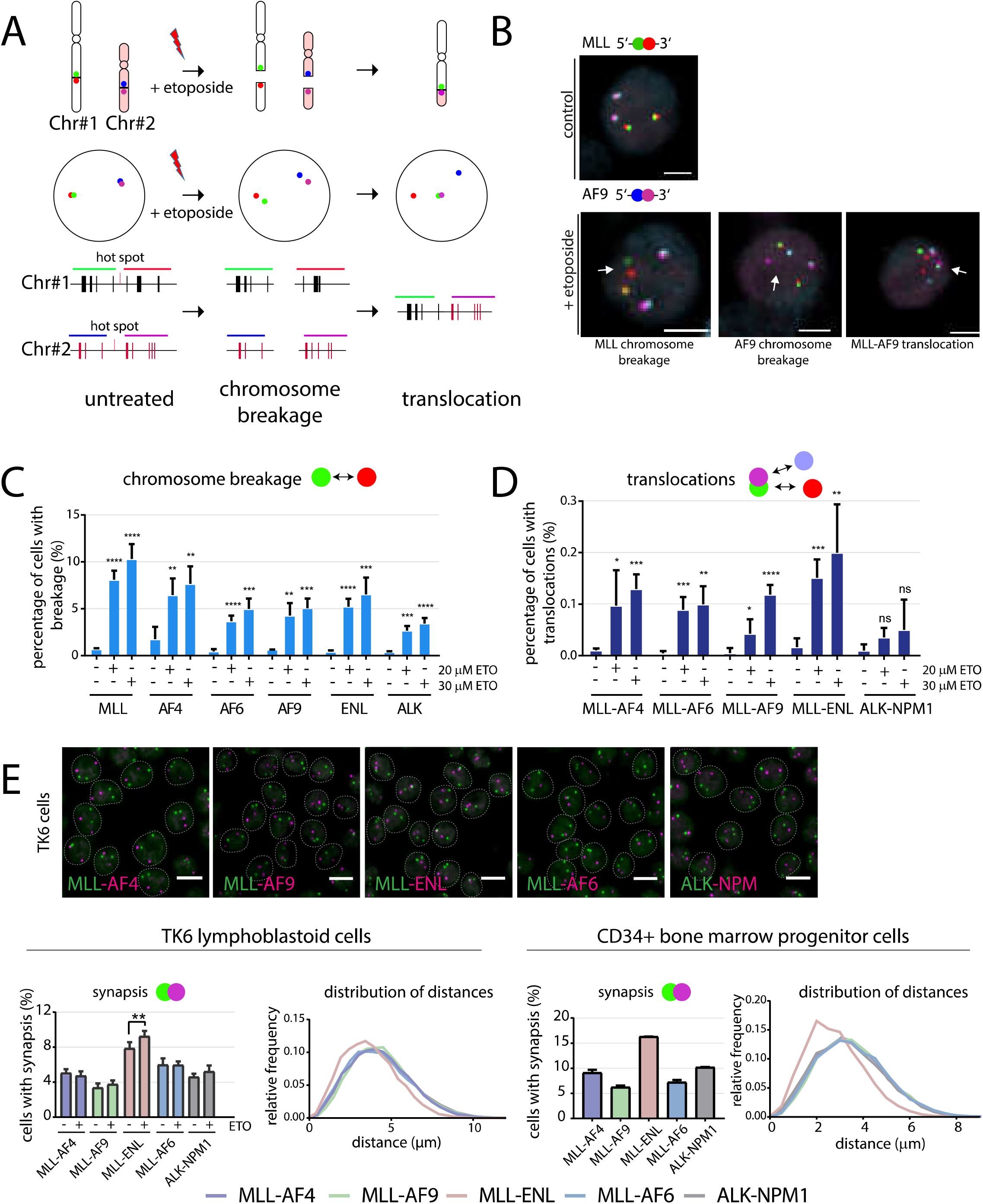
C-Fusion 3D probes ETO-induced *MLL* translocations in single cells. (A) Schematic of the methodology. (B) Representative images of untreated or released, upon ETO-treatment (30μM, 4h) TK6 cells, showing no separation, *MLL* chromosome breakage, *AF9* breakage and *MLL-AF9* translocation. Scale bar corresponds to 5μm. (C) Following release for two days upon the indicated ETO-treatment (for 4h) the percentage of cells with chromosome breakage of *MLL, AF4, AF6, AF9, ENL* and *ALK* was calculated by 4-color C-Fusion 3D in TK6 cells. Values represent means ± SD from at least four independent experiments (4000 to 16,500 cells were analyzed per sample; *P < 0.05, **P < 0.01, ***P < 0.001, ****P < 0.0001, Student t test comparison to respective untreated). (D) Frequency of cells with the indicated translocations measured by 4-color C-Fusion 3D. Cells were treated as in (C). Values represent means ± SD from at least four independent experiments (4000 to 16,500 cells were analyzed per sample; *P < 0.05, **P < 0.001, ***P < 0.0001, Student t test comparison to respective untreated). Determining the frequency of ALK-NPM1 translocations that do not occur in ETO-induced t-AML serve as control. (E) FISH images in untreated TK6 cells showing the spatial position of *MLL* (green probe) and the indicated potential partners (violet probe). Distribution of spatial distances in 3D or frequency of cells with synapsis of the indicated genes measured by C-Fusion 3D in untreated or ETO-treated TK6 cells (20μM 4h, no release) or untreated CD34+ bone marrow progenitor cells. NPM1-ALK translocations are found in cases of anaplastic large cell lymphoma (ALCL) and serve as control. Values represent means ± SD from at least two independent experiments (3700 to 6900 cells were analyzed per sample; **P < 0.01, Student t test).

Following ETO-treatment, quantification of the percentage of TK6 cells with separated *MLL* FISH probes (Figures 2A and S2A) revealed that ≈10% of the cells had experienced *MLL* breakage (9,841 events out of 98,870 cells) (Figure 2C). The frequency of cells with *MLL* breakage was cell type-specific; upon ETO-treatment (20μM, 4h) approximately 14% of K562 cells experienced *MLL* breakage (5,731 events out of 41,318 cells), while even in the presence of higher ETO-doses (60μM, 4h), only 6% of cells had the *MLL* locus broken in CD34+ hematopoietic progenitor cells (2,994 events out of 48,912 cells) (Figure S2B,C). Similar analysis showed that the most common *MLL* translocation partners break less frequently than *MLL,* with *AF4* showing slightly higher frequency (7.2% of cells, 876 events out of 12022 cells) than *AF9* (5%, 1177 events out of 23724 cells), *AF6* (5%, 714 events out of 14193 cells) and *ENL* (6.4%, 934 events out of 14532 cells) (30μM ETO, TK6 cells, Figure 2C). In contrast to the frequent breakage events, *MLL* fusions occurred rarely, at frequencies ranging from 6x10^−4^ to 2x10^−3^ in human hematopoietic cell lines and bone marrow progenitor CD34+ cells (Figures 2D and S2B,C). Interestingly, C-Fusion 3D detected *MLL-ENL* fusions two times more frequently than *MLL-AF4, MLL-AF6, MLL-AF9* (Figure 2D), although the frequency of *ENL* breakage was not significantly higher compared to the other common translocation partners (Figure 2C, p>0.05).

**Figure 3.**
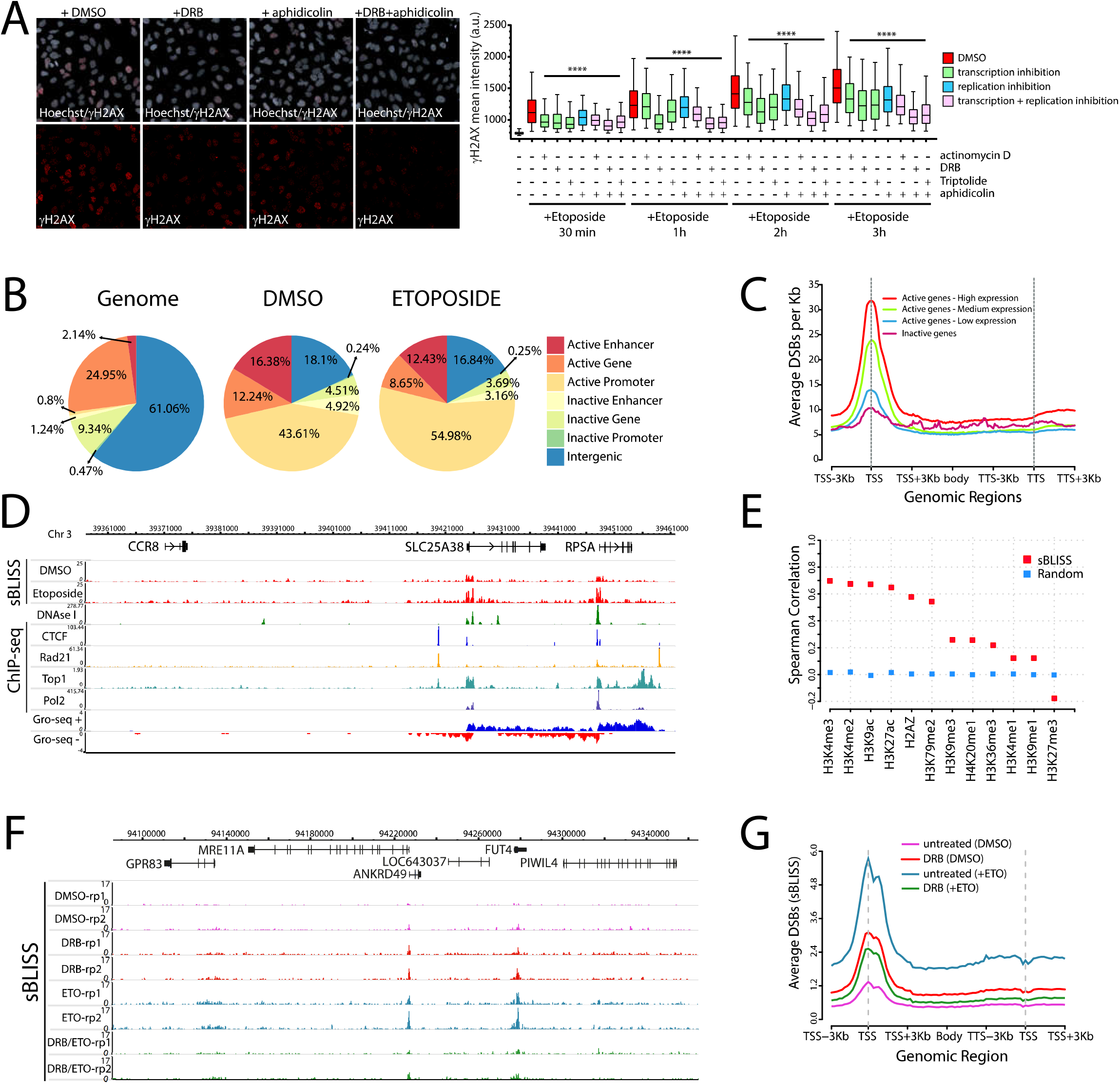
Transcription is a major contributor to TOP2-induced DNA damage. (A) Inhibition of transcription and/or replication led to a decrease in ETO-induced (10μM for indicated time) γH2AX staining in human osteosarcoma U2OS cells. Approximately 1000 cells were analyzed per condition by high-throughput microscopy and automated image analysis. Representative data and images from one out of four independent experiments are shown (****P < 0.0001, One Way Anova and Tukey test). (B) Distribution of spontaneous (DMSO) or ETO-induced (20μM, 6h) DSB-hotspots in K562 cells across genomic features. The distribution of these features in the genome is shown for comparison. (C) Aggregate plot of average DSBs per Kbp in ETO-treated K562 cells across genes classified by expression. Genes were segregated in highly expressed genes (GRO-seq RPKM count’s percentile > 90%), intermediately expressed genes (between 50% and 90%), lowly-expressed genes (< 50%) and inactive genes (< 50% and absence of H3K4me3, H3K27ac and RNA Polymerase II from promoter regions). The window spanning gene body from TSS + 3Kbp to TTS-3Kbp represents a meta-gene plot with bins regularized to span the entire gene in equal portions. (D) Genomic location showing correlation of ETO-induced DSBs by sBLISS (K562 cells) and nascent RNA expression (GRO-seq). (E) Spearman’s correlation coefficient between ETO-induced DSBs (K562 cells) and indicated histone modifications (sBLISS; red dots). As control (Random, blue), correlation of ETO-induced DSBs with randomly shuffled (one hundred times) regions of histone modifications is also shown. (F) Genomic location showing DSB profiles in TK6 cells pretreated with DRB (200μM, 3h) prior to ETO-treatment (30μM, 4h). (G) Aggregate plot of the average DSBs per Kbp across TSS, gene bodies and TTS of all genes in TK6 cells treated as in (F).

Since spatial genome organization is an important contributor to translocation frequency (Roukos and Misteli, 2014; Roukos et al., 2013; Zhang et al., 2012), we asked whether the higher frequency of *MLL-ENL* translocations is merely a consequence of the closer proximity between the *MLL* and the *ENL* loci in the three dimensional (3D) nuclear space. To assess this directly, we measured the Euclidean 3D distances between *MLL* and the most frequent translocation partner in untreated or ETO-treated TK6 cells. In line with the observed higher frequencies of *MLL-ENL* translocations, *MLL* and *ENL* genes were closer in proximity compared to other *MLL* fusion partners (Figure 2E, distribution of distances) and *MLL* and *ENL* alleles were synapsed more frequently compared to other fusion partners (Figure 2E; MLL-ENL=7.9%, MLL-AF4=5.1%, MLL-AF6=6.0%, *MLL*-AF9=3.4%, *ALK-NPM1=4.6*% *p*<0.01). The spatial arrangement of these genes was conserved in the CD34+ bone marrow progenitor cells (Figure 2E), in which ETO-treatment resulted in an additional increase in synapsis of the *MLL* gene with *ENL,* but not with other partners (Figure 2E, *p=0.004).* Thus, upon ETO-treatment, *MLL* translocations occur at frequencies ranging from 6x10^−4^ to 2x10^−3^ in hematopoietic cell lines and progenitor cells and the apparent higher spatial proximity of *MLL* with *ENL* favors the formation of *MLL-ENL* fusions.

### Transcription is a major contributor to TOP2-induced DNA damage

To explore the contribution of active transcription and replication to the conversion of TOP2ccs to DSBs, we pre-treated cells with a variety of transcription and replication inhibitors and quantified the activation of the DNA damage response (DDR) upon ETO treatment across the cell cycle by high-throughput imaging. Pre-treatment of cells with transcription inhibitors, which either prevent transcriptional initiation or elongation (Bensaude, 2011), led to a substantial decrease of ETO-induced DNA damage (marked by γH2AX levels) in all cell cycle phases (Figures 3A, S3A and S3B, *p*<10^−4^). Similarly, inhibition of replication (Figure S3A), led to a decrease in ETO-induced γH2AX and ETO-induced replication stress in S phase (marked by pRPA S4/S8 levels) (Figure S3B,C *p*<10^−4^), while co-inhibition of both replication and transcription had a synergistic effect (Figure 3A). These data support the notion that both active transcription and replication contribute to the conversion of TOP2ccs to DSBs.

To further evaluate the contribution of transcription to the conversion of TOP2ccs to DSBs, we assessed the genome-wide distribution of ETO-induced DSBs relative to the transcriptional status. More than 75% of the ETO-induced hot spots in K562 cells were localized within transcriptionally active regions of the genome (active promoters, active enhancers, active genes, Figure 3B), while only 7% of them were found within inactive regions. Notably, similar enrichment of DSBs in active regions was observed in cells non-treated with ETO (Figure 3B, DMSO), indicating that ETO enhances the frequency of breakage in active regions that are, *per se,* more fragile compared to the rest of the genome. Additionally, genome-wide comparison of transcriptional output measured by global run-on sequencing (GRO-seq) with ETO-induced breakage measured by sBLISS, revealed a strong correlation (Spearman’s correlation=0.64, Figure S3D, p<10^−5^). In line, highly transcribed genes accumulated more ETO-induced DSBs at their promoters and gene bodies compared to genes that were moderately transcribed, transcribed at low levels or were inactive (Figures 3C and 3D). A similar trend was observed in untreated K562 cells for endogenous DSBs (DMSO, Figures 3B and S3E). To corroborate these observations, we correlated ETO-induced DSB-hot spots with a variety of histone marks assessed by ChIP-seq. We found that DSB-hot spots were enriched within chromatin regions marked by H3K4me3, H3K9ac, H3K27ac (Spearman’s correlation: 0.69 [*p*<10^−5^], 0.67 [*p*<10^−5^], 0.64 [*p*<10^−4^], respectively), but were less enriched or depleted in regions with histone marks representing heterochromatin and/or suppressed transcription, such as H3K9me3 and H3K27me3 (*p*<10^−3^) (Figures 3E and S3F).

To further understand the relationship between ETO-induced DSBs and transcription, we blocked RNA polymerase II elongation with 5,6-dichloro-1-beta-D-ribofuranosyl-benzimidazole (DRB), before treatment of cells with ETO, and we then, profiled DSBs across the genome by sBLISS. Similar to the observed reduction of ETO-induced DDR upon inhibition of transcription (Figures 3A and S3B), pre-treatment with DRB led to a global decrease in ETO-induced DSBs (Figure 3F), a reduction which was apparent both across gene bodies and promoter regions (Figure 3G). Overall, our data show that ETO-induced DSBs are highly enriched within active genomic regions and are dependent on elongation of RNA polymerase II, demonstrating that transcription is a major contributor to the occurrence of TOP2-induced DSBs.

### TOP2-induced DSBs are enriched at loop anchors and highly correlate with transcriptional output and directionality

Given that TOP2B, cohesin and CTCF interact and are found enriched at chromatin loop anchors and topological domain borders (Uuskula-Reimand et al., 2016), as well as that TOP2-induced DSBs are largely transcription-dependent (Figures 3 and S3), we sought to explore the relationship between TOP2-induced breakage at CTCF binding sites within loop anchors, while also considering the contribution of transcription in this process. Expression analysis revealed that *MLL* and the vast majority of recurrently identified fusion partner genes (known to contribute to acute leukemias, *MLL* recombinome, ≈89 genes (Meyer et al., 2018), were amongst the most highly expressed genes in human hematopoietic cell lines, as well as in human hematopoietic CD34+ stem cells (Figures 4A and S4A). Strikingly, we found that *MLL* and the potential translocation partner genes are highly enriched at, or close to, chromatin loop anchors across various human hematopoietic cell lines, and other cell types (Figures 4B and S4B, and data not shown). These findings suggest the intriguing possibility that high transcriptional activity at loop anchors might require TOP2 functions, which in turn intrinsically bears the potential of contributing to genomic instability favoring the formation of oncogenic translocations. To address this, we asked whether there is a correlation between transcriptional output, TOP2-induced chromosome breakage, and localization at loop boundaries. Indeed, in addition to the observed positive correlation between ETO-induced DNA damage and transcriptional output (Figures S3D and 4C), we can show that gene expression inversely correlates with the genomic distance of the gene from the closest loop anchor (Figure S4C, TK6 cells: Spearman’s correlation=-0.83; K562 cells: Spearman’s correlation = −0.81). Likewise, we found that TOP2-induced DSBs levels within a given gene inversely correlated with the distance to the closest loop anchor (Figure S4C, TK6 cells: Spearman’s correlation=-0.80; K562 cells: Spearman’s correlation=-0.78), indicating that genes with higher fragility due to TOP2 activity tend to be more proximal to loop anchors. Strikingly, increased chromosome fragility together with localization at such insulated boundaries were strongly linked to high gene expression, and *MLL* and several fusion partner genes shared all three features (Figure 4C, TK6 cells: triple correlation=0.618; K562 cells: triple correlation=0.623, *MLL* and fusion partners marked as blue circles). This triple association was lost when we performed cross-cell line correlations (Figure S4D), indicating that high transcription, localization at chromatin loop boundaries, and DNA fragility, identified features of *MLL* and fusion partner genes, are to a large extend cell type-specific and inherently linked.

**Figure 4.**
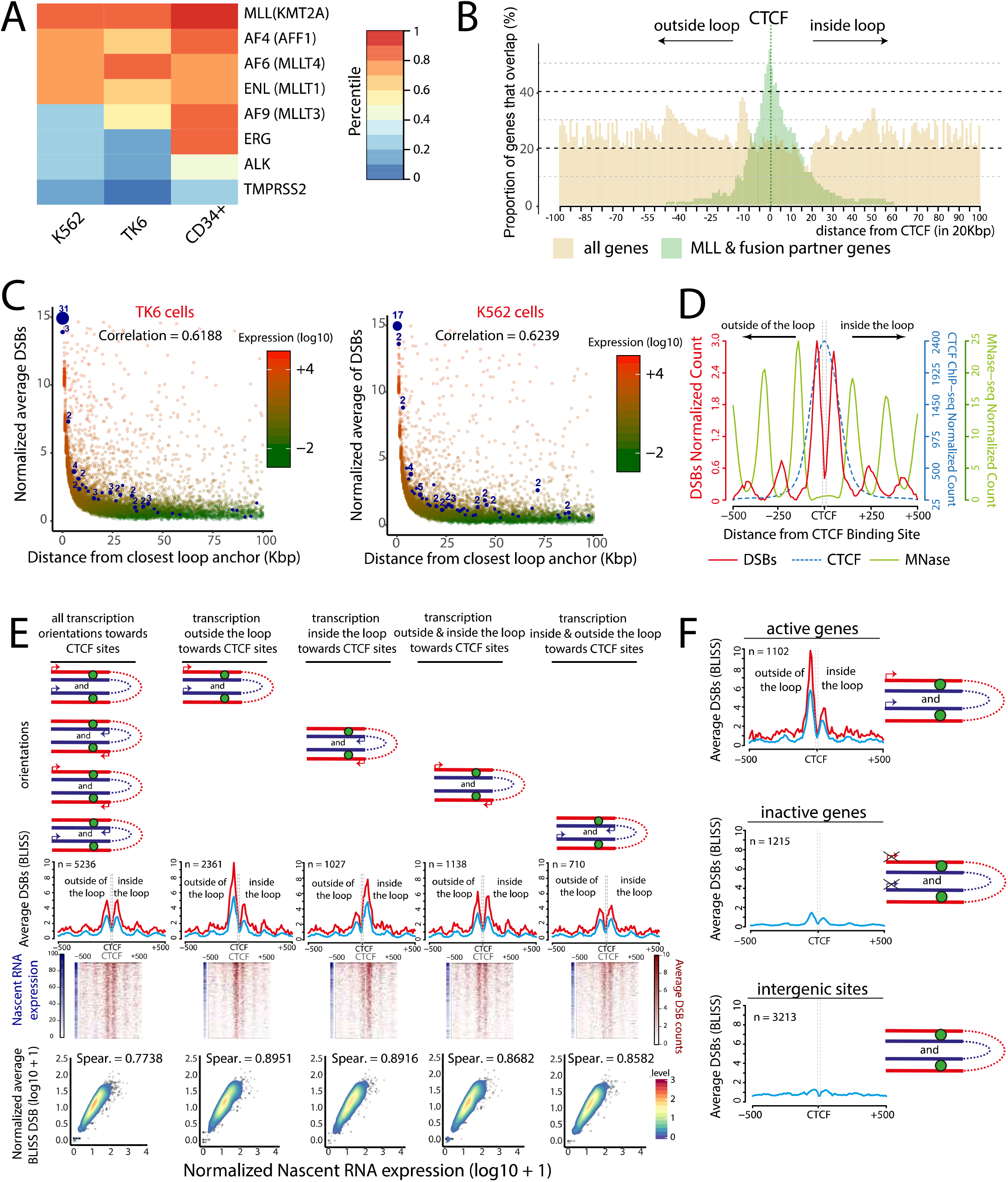
TOP2-induced DSBs are enriched at loop anchors and highly correlate with transcription output and directionality. (A) Heatmap showing expression values of the *MLL* and fusion partner genes *(AF4, AF6, ENL, AF9)* and genes that recurrently translocate in prostate cancer *(ERG, TMPRSS2)* (Haffner et al., 2010) or anaplastic large cell lymphoma *(ALK)* (Roukos and Mathas, 2015), given as percentiles of the genome wide expression levels in the hematopoietic K562 and TK6 cells and the CD34+ stem cells. (B) Histogram of the proportion of overlapping gene bodies of *MLL* and fusion partners genes (green) or control genes (all genes in the genome; yellow), relative to loop anchor sites in TK6 cells. Genes were placed with regard to their respective distance between the closest loop anchor’s CTCF position and the gene’s TSS. (C) Correlation of ETO-induced DSBs of genes as a function of distance from their closest loop anchor, and transcription output levels, in TK6 and K562 cells. Distances were rounded up to the nearest Kbp and were measured from the closest gene end (TSS or TTS) to the middle of the closest loop anchor; distances of 0 bp correspond to an overlap between the gene and the 10-Kbp loop anchor. *MLL* and fusion partner genes represented by blue circles with size corresponding to the number of overlapping genes (numbers shown on top). (D) Aggregate plot of ETO-induced DSBs (K562 cells, red line) centered around (+/-500bp) the CTCF motives (dashed grey lines) within loop anchors, and superimposed nucleosome occupancy (MNase-seq, green line) and CTCF binding (ChIP-seq signal, blue dashed line). (E) Average ETO-induced DSBs (TK6 cells, sBLISS) at intragenic loop anchor sites were plotted around +/-500bp from CTCF motives (dashed grey lines, CTCF sites are depicted at loop structures as green circles) according to transcription orientation (red and blue arrows, represent transcription direction of genes in the forward and reverse strands, respectively). Aggregate plots of average DSBs (upper panel) and heatmaps (middle panel) around the CTCF motives are shown (blue heatmaps depict nascent RNA expression at these sites). Blue lines in aggregate plots represent DSB signals of all genes at the region and red lines DSB-signals of genes with expression higher than the 90% percentile. Scatterplots (and their respective Spearman’s correlation coefficients) between sBLISS DSB-counts and nascent RNA expression at these sites (GRO-seq) are also shown (lower panel). The color key of the correlation scatterplots represents a twodimensional kernel density estimation. Genes intersecting with the downstream anchor of their respective loop were mirrored with regard to the center of the plot (CTCF position). (F) Aggregate plots of ETO-induced DSBs (K562 cells, sBLISS) at CTCF sites found at loop anchors within active genes, inactive genes or intergenic regions. The plot follows the same rationale as the aggregate plots in (E).

In agreement with these findings, ChIA-PET data from CTCF and cohesin subunit (Rad21) interactions in human hematopoietic cells showed that the *MLL* translocation hot spot localizes at a loop anchor region establishing loops with upstream genomic regions (Figure S5A). In line with our observations above, ETO-induced DSBs were enriched within the loop and 5’ to the CTCF motif (Figure S5A, inset), suggesting that TOP2 induced breakage near the CTCF site of the translocation hotspot correlates with the directionality of transcription. To investigate this further, we probed the distribution of ETO-induced DSBs relative to the center of the G-rich, oriented, CTCF binding motif found at loop anchors across the genome. In accordance with what was previously reported for mouse B-lymphocytes (Canela et al., 2017), TOP2-induced DSBs were enriched at nucleosome free regions around the strongly positioned nucleosomes of CTCF sites (Figure 4D, comparison to MNase-seq). ETO-induced DSB sites showed two peaks relative to the CTCF binding motif, both at a mean distance of 45 nucleotides from the CTCF motif, indicating that TOP2-induced breaks occur in both directions, inside and outside, of the loops (Figure 4D). To investigate whether TOP2-induced breakage at CTCF sites shows any directionality bias, we generated aggregate plots of ETO-induced breakage at CTCF sites within genes at loop anchors classified according to their transcriptional orientation (Figure 4E). Strikingly, the orientation of transcription dictated a polarized TOP2-induced breakage pattern around CTCF sites, with a prominent enrichment of TOP2-induced DSBs between the direction of transcription and the CTCF binding site (Figure 4E). When transcription was directed towards the CTCF sites from genomic regions outside the loop, TOP2-induced DSBs were found enriched at a mean distance of 45 nt just outside the loop. In contrast, when transcription was directed towards the CTCF anchor sites from regions within the chromatin loops, TOP2-induced breakage was enriched at a similar distance, but on the inside of the chromatin loop (Figure 4E). TOP2-induced breakage at CTCF sites within loop anchors highly correlated with transcriptional output (Figure 4E, Spearman’s correlation≈0.89) and was greatly reduced for inactive genes or non-transcribed intergenic sites, indicating that it is predominantly transcription-dependent (Figure 4F). Similar to *MLL* (Figure S5A), TOP2-induced DSBs at fusion partner genes located at loop anchors were also asymmetric and positioning of breakage again correlated with transcriptional directionality (Figure S5B). Taken together, our findings demonstrate that TOP2-induced DNA fragility, transcriptional activity and localization at the bases of chromatin loops go hand-in-hand, and break patterns at highly transcribed *MLL* and fusion partner genes located at loop anchors, are asymmetric and strongly linked to transcriptional output and directionality.

### *MLL* translocations are largely transcription-dependent

Given that the *MLL* and several fusion partner genes are located in the vicinity of loop boundaries and are highly transcribed, our findings support the intriguing possibility that transcription-induced torsional stress towards stabilized loop anchors, bound by CTCF and cohesin, is released by directed TOP2 activity between the CTCF sites and the direction of transcription. Under conditions of incomplete TOP2 activity, such as in the presence of ETO, unresolved complexes are converted to DSBs that promote the formation of *MLL* fusions. To test this possibility, we directly assessed the contribution of transcription to the susceptibility of *MLL* and the recurrent partner genes to breakage and the formation of *MLL* fusions. TK6 cells pretreated with transcription or replication inhibitors, were exposed to ETO, and the percentage of cells with breakage at *MLL* or at the recurrent translocation partners was assessed by C-Fusion 3D. Replication inhibition slightly reduced the percentage of cells with *MLL* or *AF4* breakage (Figure 5A), whereas transcription inhibition resulted in an even more significant reduction of the percentage of cells with chromosome breakage within *MLL* and all potential translocation partners *(MLL* from 9.7% to 3.4%, *p*<10^−3^; *AF4, AF9, ENL, AF6* by two-fold *p*< 0.05), (Figure 5A). Importantly, inhibition of transcription elongation by DRB, also reduced the frequency of *MLL* fusions with the various partners to almost background levels (Figure 5B), suggesting that active transcription is the predominant activity that contributes to therapy-related ETO-induced *MLL* fusions.

**Figure 5.**
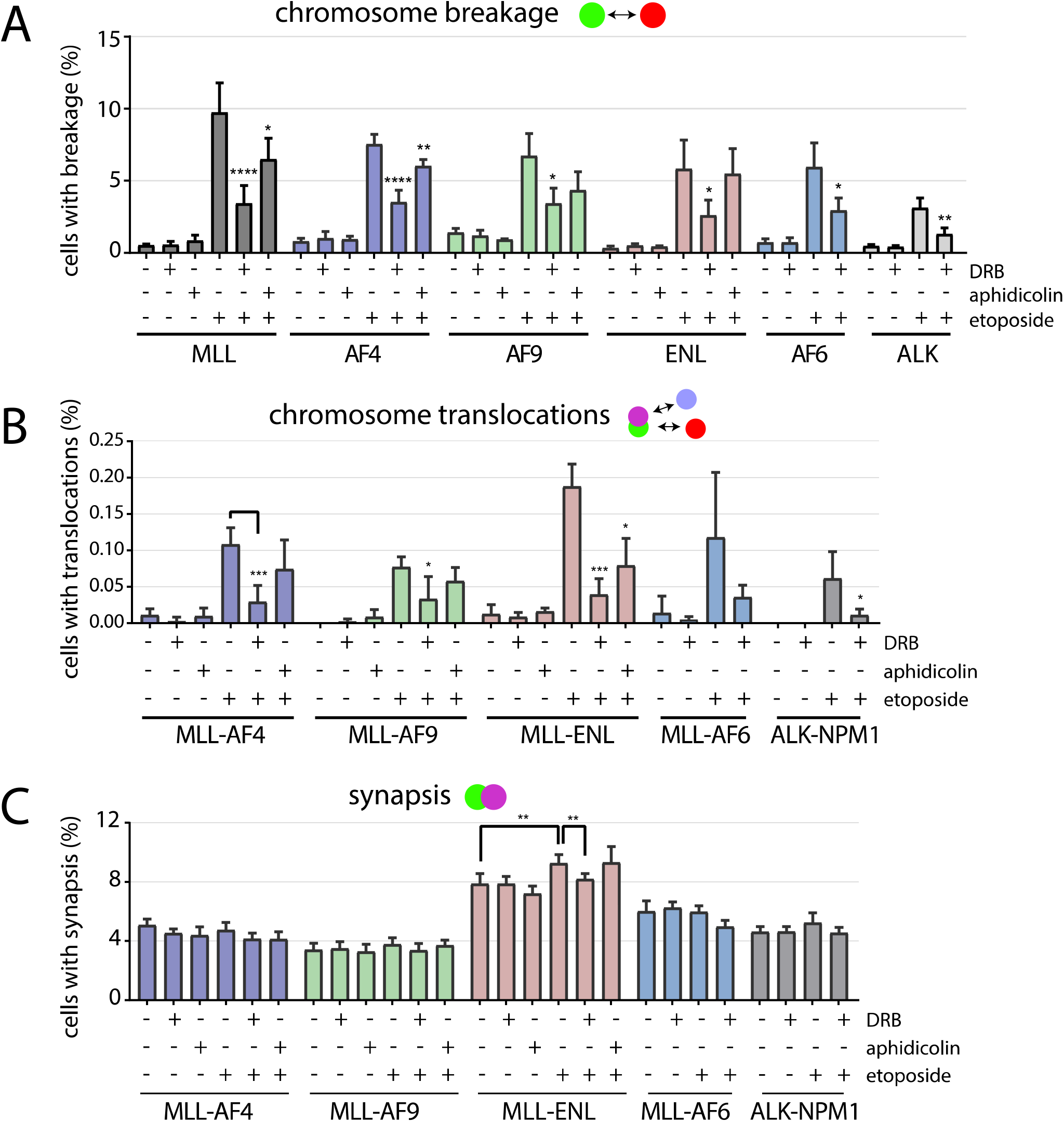
The formation of *MLL* fusions is largely transcription-dependent. (A) TK6 cells were pretreated with DRB (200μM, 3h) or aphidicolin (8μM, 3h), prior to the treatment with ETO (20μM, 4h). Two days upon release from ETO and inhibitors, the percentage of cells with chromosome breakage of *MLL, AF4, AF6, AF9, ENL, ALK* (A) and indicated translocations (B) was calculated by 4-color C-Fusion 3D. Values represent means ± SD from at least three independent experiments (4000 to 16,500 cells were analyzed per sample; *P < 0.05, **P < 0.01, ***P < 0.001 ****P < 0.0001, Student t test comparison to ETO-treatment). (C) Frequency of cells with synapsis of the indicated genes. Cells treated as in (A). Values represent means ± SD from at least three independent experiments (7500 to 16,500 cells were analyzed per sample; **P < 0.01 Student t test comparison as indicated).

It has been previously proposed that transcription may contribute to the formation of *MLL* translocations by keeping *MLL* and its recurrent translocation partner genes in close proximity, within common transcription factories (Cowell et al., 2012). To assess whether the observed effect of transcription inhibition in reducing *MLL* fusion is due to an influence on the spatial proximity of *MLL* and potential translocations partners genes, we calculated the 3D Euclidean distances between *MLL* and the recurrent fusion partners upon transcriptional inhibition. In absence of DNA damage, inhibition of transcription by DRB, did not influence the percentage of cells with synapsed *MLL* and partner genes (Figure 5C), suggesting that transcription does not contribute to the spatial proximity between *MLL* and the fusion partners. However, upon ETO-treatment, the fraction of cells with synapsed *MLL* and *ENL* genes, which intriguingly were in closer proximity compared to the other fusion partners in both TK6 and CD34+ stem cells (Figure 2E), was increased (Figure 5C). The observed increase in synapsis was sensitive to transcriptional inhibition (Figure 5C), suggesting that transcription-dependent DNA damage, rather than a direct role of transcription in inducing proximity of the translocation partners, favors the formation of *MLL* translocations. Collectively, our data suggest that transcription-dependent induction of TOP2-mediated DSBs at highly transcribed genes at loop anchors drive the formation of *MLL* fusions.

### Etoposide-induced DSBs and *MLL* translocations depend on the activity of both TOP2 isoforms

ETO-induced DNA damage across the genome (Azarova et al., 2010; Azarova et al., 2007; Canela et al., 2017) or at the *MLL* locus (Canela et al., 2017; Cowell et al., 2012) has been attributed mainly to functions of the TOP2B isoform, but exclusive TOP2A isoform-dependent functions have also been reported (de Campos-Nebel et al., 2010; Tammaro et al., 2013). To systematically explore the contribution of both TOP2 isoforms to ETO-induced DSBs and the formation of *MLL* fusions, we engineered a colorectal carcinoma HCT116 cell line in which both endogenous *TOP2A* alleles, are tagged with the auxin-inducible degron (AID) (Natsume et al., 2016). The resulting TOP2A-AID cell line allows to rapidly eliminate up to 95% of the endogenous TOP2A pool upon addition of auxin (Figure 6A). In addition, we used CRISPR/Cas9 technology to target the *TOP2B* alleles and generate isogenic cell lines in which each TOP2 isoform separately, or both isoforms together, can be efficiently knocked out (Figure 6A). Following ETO-treatment, quantification of DDR activation by γH2AX phosphorylation revealed that both TOP2A and TOP2B contribute to ETO-induced DNA damage and elimination of both isoforms led to a further decrease of ETO-induced DSBs (Figure S6A). In line with this observation, breakage at the *MLL* locus was also dependent on both isoforms, although elimination of TOP2A showed a more pronounced effect (Figure 6A). To validate our findings in other cell types, we quantified ETO-induced DNA damage in the previously characterized human fibrosarcoma cell line HT1080, in which the highly expressed TOP2A isoform (compared to TOP2B expression), can be eliminated upon doxycycline treatment (Carpenter and Porter, 2004) (Figure S6B). In these cells, ETO-induced DNA damage (Figure S6B), *MLL* breakage and *MLL* translocations (Figure 6B), were exclusively TOP2A-dependent, suggesting that the relative expression levels of the TOP2A and TOP2B isoforms, might be predictive of the isoform-specific dependency to ETO-induced DNA damage. Indeed, probing the relative expression of the TOP2 isoforms and the isoform-specific dependency to ETO-induced DNA damage in a panel of different cell lines (Figures S6C-E), showed that ETO-induced DNA damage depends on both isoforms. In support, *MLL* breakage and *MLL-AF4* translocations were dependent on both isoforms in the breast carcinoma cell line Cal51, which expresses both isoforms at average levels (Figures 6C and S6D) and in CD34+ human hematopoietic progenitor cells (Figure 6D), suggesting that trapping of either of TOP2A or TOP2B isozymes is responsible for the occurrence of *MLL* fusions.

**Figure 6.**
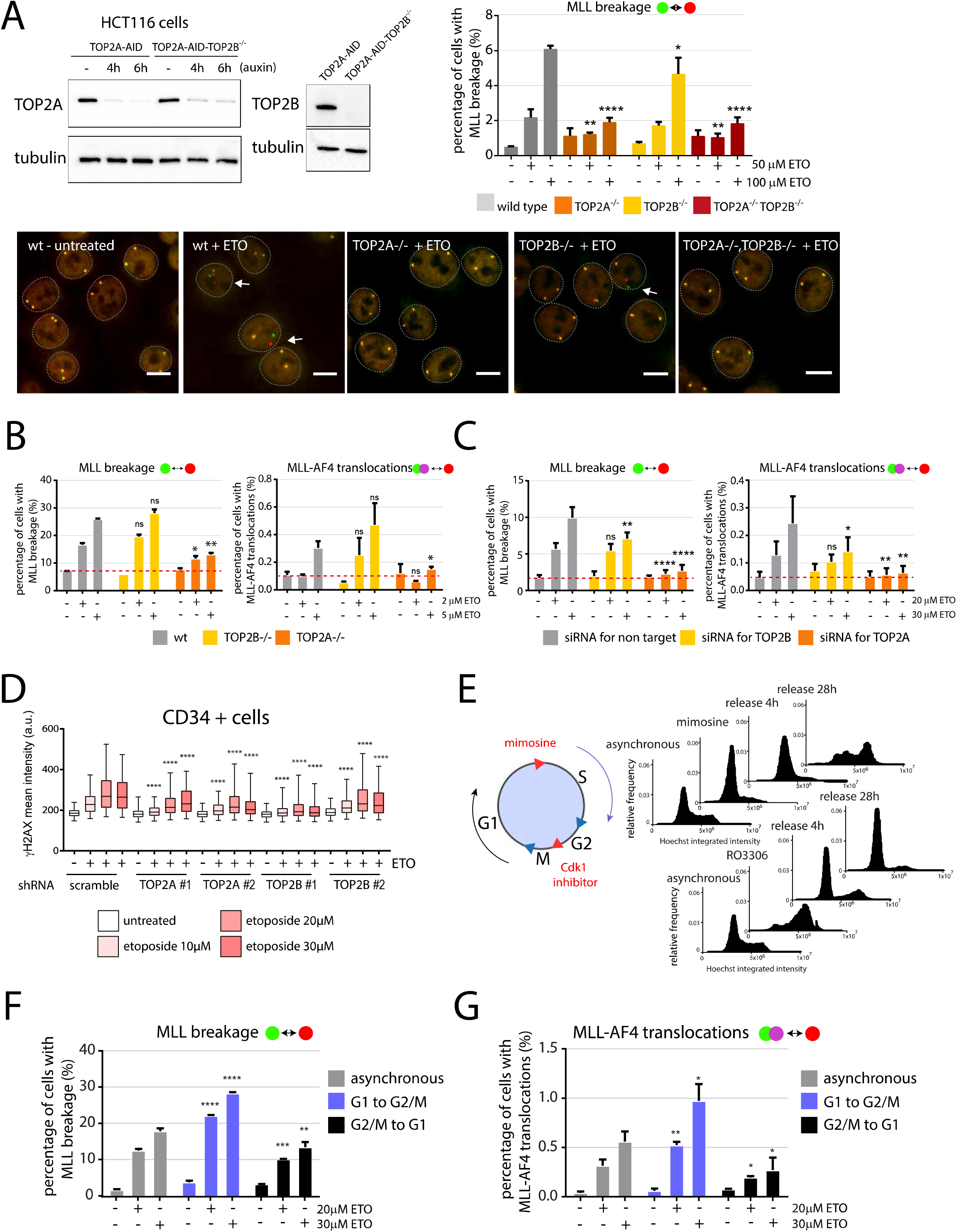
Both TOP2 isoforms contribute to ETO-induced DSBs and *MLL* translocations. (A) Verification of the engineered TOP2A-mAID and TOP2A-mAID-TOP2B^−/-^ HCT116 cells by assessing TOP2 isoform-specific levels upon auxin treatment by western blotting (left panel). Right panel: percentage of HCT116 cells with *MLL* breakage upon ETO treatment (50μM or 100 μM for 3h) assessed by C-Fusion 3D upon depletion of TOP2A by auxin (500μM, 4h pretreatment) or in cells deficient for TOP2B (1700 to 7300 cells were analyzed per sample in four independent experiments, means ± SD). Representative images of cells upon indicated conditions is shown (lower panel, green probe is *MLL* 5’, red probe is *MLL* 3’ see also Figure S2A, scale bar equal to 10μm). (B) Percentage of HT1080 cells deficient for TOP2A or TOP2B with *MLL* breakage (left) and *MLL-AF4* translocations (right) upon ETO (2μM or 5μM for 4h, 24h release) assessed by C-Fusion 3D (1500 to 5600 cells were analyzed per sample, two independent experiments, means ± SD). (C) Both TOP2A and TOP2B isoforms contribute to ETO-induced *MLL* breakage and *MLL* translocations in human Cal51 cells upon silencing of the respective isoforms by siRNA an ETO (20μM or 30μM for 4h, 24h release) (1500 to 8100 cells were analyzed per sample, three independent experiments, means ± SD). (D) ETO-induced DSBs in CD34+ progenitor cells are both TOP2A and TOP2B-dependent. CD34+ cells were transduced with different shRNAs for the two TOP2 isoforms and ETO-induced DSBs were quantified by γH2AX staining upon ETO treatment (4h) at the indicated doses. GFP positive cells were selected by image analysis; at least 700 GFP positive cells per condition were quantified for γH2AX mean intensity. Representative box plot of two independent experiments is shown. Statistical significance was calculated with One Way ANOVA test and Tukey compared to Scramble. (E) TK6 cells were arrested in G1 or G2/M phases by treatment with mimosine or the Cdk1 inhibitor (RO3306) respectively, and were released into the next cell cycle phase (S and G1, respectively) in the presence of ETO (20 or 30μM for 4h). After ETO treatment, cells were maintained in the second inhibitor for 24h. Cell cycle distribution at indicated time-points was assessed by measuring DNA content from Hoechst staining (Roukos et al., 2015). (F) Cell cycle regulation of *MLL* breakage and (G) *MLL-AF4* translocations. Cells were treated as in (E) and upon release from ETO for one day (in the presence of RO3306 or mimosine, prohibiting cells to leave G2/M or G1, respectively), the percentage of cells with *MLL* breakage and *MLL-AF4* translocations was assessed by C-Fusion 3D (1500 to 14,500 cells were analyzed per sample, three experiments, means ± SD). (A-G) *P < 0.05, **P < 0.01, ***P < 0.001, ****P < 0.0001, Student’s t test comparison to respective treatment in wild type cells (A to C) or to asynchronous cells (F, G).

Given that the TOP2A expression is regulated through the cell cycle (Pommier et al., 2016), we addressed whether the occurrence of ETO-induced *MLL* breakage and *MLL* translocations is cell cycle phase-dependent. Cells arrested in G1 by the use of mimosine, were released in S phase in the presence of ETO. Alternatively, cells arrested in late G2 by the use of the Cdk1 inhibitor (RO3306) were released in G1 in presence of ETO (Figures 6E and S6F). Twenty four hours after release from ETO-treatment, the percentage of cells with detected *MLL* breakage or *MLL-AF4* fusions was determined by C-Fusion 3D. Our analysis revealed a two-fold increase in the population of cells with *MLL* breakage and translocations in S/G2 phases, where both TOP2 isoforms are highly expressed, compared to G1, where the TOP2B form is predominantly expressed (Heck et al., 1988; Hsiang et al., 1988; Woessner et al., 1991) (Figure 6F,G). Altogether, these results demonstrate that both TOP2 isoforms contribute to *MLL* breakage and *MLL* fusions.

### MRE11, TDP2 and NHEJ suppress *MLL* translocations

Trapped topoisomerases on DNA 5’ termini are removed by the phosphodiesterase activity of tyrosyl-DNA phosphodiesterase 2 (TDP2) that cleaves the bond between the topoisomerase and the 5’ phosphate of the DNA, generating ends that can be readily ligated by the non-homologous end-joining (NHEJ) pathway (Cortes Ledesma et al., 2009; Gomez-Herreros et al., 2013). Alternatively, MRE11, as part of the MRN complex (MRE11/RAD50/NBS1), can nucleolytically remove TOP2-DNA adducts, leaving DNA ends which can be channeled to either NHEJ or homologous recombination (HR) (Hoa et al., 2016; Stingele et al., 2017). How factors involved in the removal of TOP2-DNA covalent complexes and of the downstream DSB-repair pathways contribute to *MLL* breakage and the formation of oncogenic *MLL* translocations remains poorly understood.

To address the contribution of factors involved in the removal of TOP2-DNA covalent complexes to ETO-induced breakage and the formation of *MLL* translocations, we employed a panel of isogenic human lymphoblastoid TK6 cell lines conditionally disrupted for Mre11 (MRE11^−/-^), its nuclease-activity (MRE11^−/H129N^) (Hoa et al., 2016), the resection protein CtIP (CtIP^−/-^) (Hoa et al., 2015), the phosphodiesterase TDP2 (TDP2V^−/-^) and combinations thereof (Figures 7 and S7B). In line with previous findings showing that TDP2 protects cells from TOP2-induced DSBs (Gomez-Herreros et al., 2013; Gomez-Herreros et al., 2017), TDP2 deficient cells show impaired capacity in repairing ETO-induced DSBs (Figure S7A). Delayed repair kinetics upon ETO-induced DNA damage, were also evident in cells deficient for NHEJ (cells deficient in LIG4^−/-^), but not for HR (cells deficient in RAD54 or CtIP) (Figure S7A), suggesting that TOP2-inducedDSBs are predominantly repaired by NHEJ (Pommier et al., 2016). In contrast to TDP2 deficiency, cells deficient in MRE11 nuclease activity or double MRE11/TDP2 null cells (MRE11^−/H129N^/TDP2^−/-^), showed indistinguishable repair kinetics compared to control cells upon release from ETO-treatment. Since MRE11 nuclease activity is essential to remove TOP2-DNA covalent complexes (Deshpande et al., 2016; Hoa et al., 2016), this effect is probably a reflection of abrogated signaling towards H2AX in absence of MRE11 nuclease activity.

**Figure 7.**
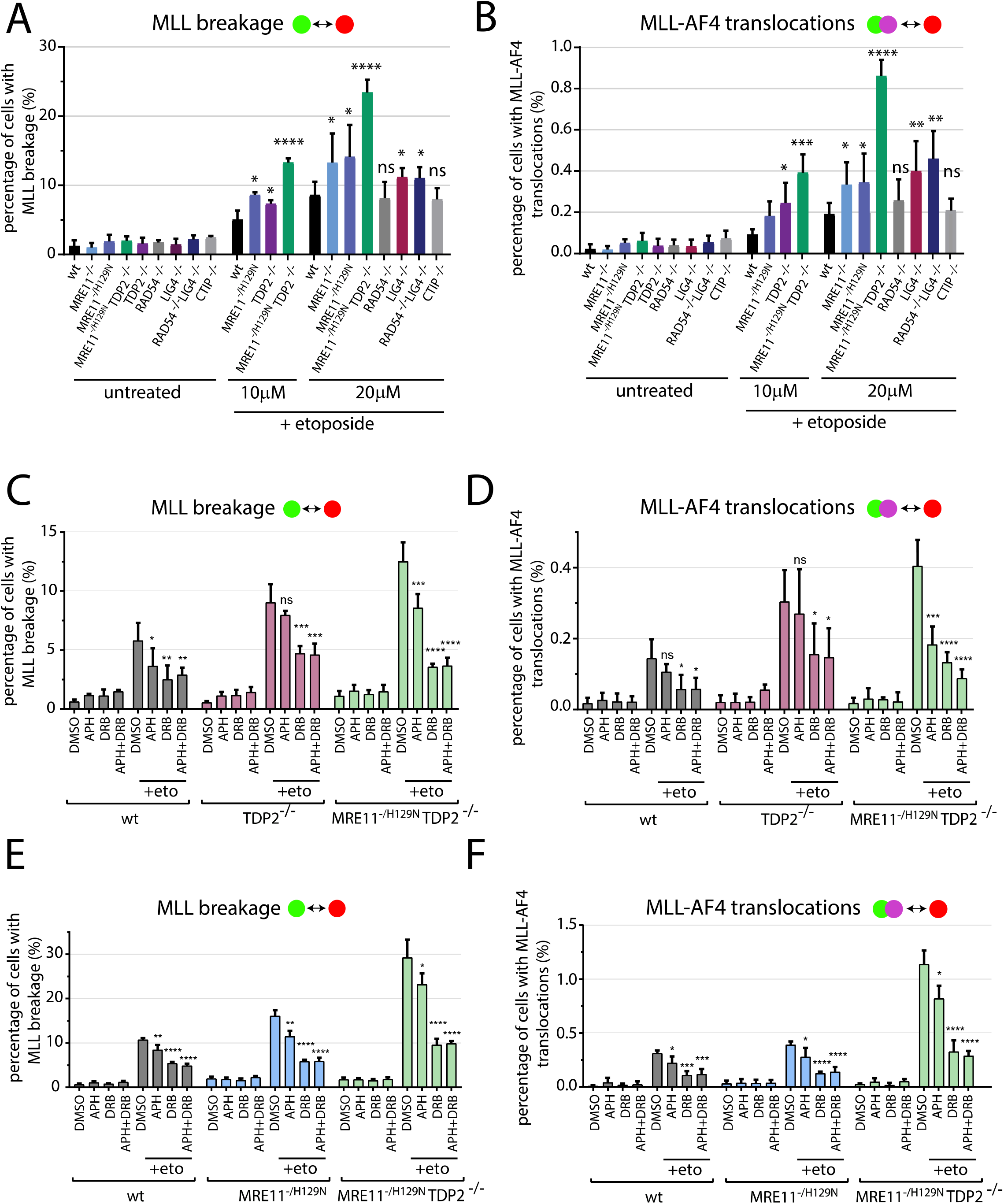
TDP2, MRE11 and NHEJ suppress the formation of *MLL* translocations. (A), (B) TK6 cells, deficient for indicated genes, were treated with 10 or 20μM ETO for 4h and released for two days. The frequencies of *MLL* breakage (A) and *MLL-AF4* translocations (B) were assessed by C-Fusion 3D. Values represent means ± SD from at least four independent experiments (1500 to 16,000 cells were analyzed per sample). (C)-(F), Parental, TDP2^−/-^, MRE11^−/H129N^ or double mutant TDP2^−/-^/MRE11^−/H129N^ TK6 cells were treated with 200μM DRB, 8μM aphidicolin or both for 3 hours, prior to 10μM or 20μM ETO-treatment for 4h. The frequencies of *MLL* breakage (C, E) and *MLL-AF4* translocations (D, F) were assessed by C-Fusion 3D two days after release from ETO treatment. Values represent means ± SD from at least four independent experiments (1800 to 16,500 cells were analyzed per sample). *P < 0.05, **P < 0.01, ***P < 0.001, ****P < 0.0001, Student t test comparison to respective treatment in wild type cells (A, B) or to DMSO treated cells (C-F).

We then assessed the contribution of these factors to the susceptibility of the *MLL* locus to breakage and the formation of *MLL* fusions by employing C-Fusion 3D upon release from ETO-treatment. Compared to parental cells, deficiency in HR showed no effect in the frequency of *MLL* breakage and the formation of *MLL-AF4* translocations, in line with the limited functions of HR in repairing ETO-induced DSBs (Figure 7A,B, p>0.05) (Ashour et al., 2015; Pommier et al., 2016). Deficiency in LIG4^−/-^ or in both LIG4^−/-^ and RAD54^−/-^, led, however, to a two-fold increase in frequency of cells with *MLL* breakage and *MLL-AF4* translocations upon ETO-treatment (Figure 7A,B). These data suggest that DSB repair by NHEJ suppresses the occurrence of *MLL* fusions by favoring intrachromosomal repair of TOP2-generated DSBs. A similar increase in the percentage of cells with *MLL* breakage and *MLL-AF4* translocations was observed upon ETO-treatment in cells deficient for TDP2 or MRE11 nuclease activity (Figure 7A,B). These findings indicate that TDP2 and the nuclease activity of MRE11 act as suppressors of *MLL* translocations by removing prone to breakage TOP2-DNA covalent complexes upon ETO-treatment within the *MLL* gene. Importantly, deficiency for both TDP2 and the nuclease activity of MRE11 led to a further increase in *MLL-AF4* translocation frequency (*p*<10^−3^, Figure 7B, four-fold increase over control), revealing a non-epistatic relationship between these factors in suppressing the occurrence of *MLL* fusions.

Since ETO-induced TOP2-DNA covalent complexes can be converted to DSBs by interference of the replication machinery and/or transcription elongation, we sought to examine whether the functions of TDP2 and MRE11 in suppressing *MLL* fusions were specifically associated with TOP2-induced DSBs arising during these processes. To this end, we pretreated parental, TDP2^−/-^, MRE11^−/H129N^ or double deficient (TDP2^−/-^/MRE11^−/H129N^) TK6 cells with DRB and/or aphidicolin, and upon release from ETO treatment we calculated the frequency of cells with *MLL* breakage and *MLL-AF4* fusions. Inhibition of transcriptional elongation, but not of replication, led to a great reduction in *MLL* breakage and translocations in TDP2 null cells, suggesting that TDP2 performs a protective role in preventing TOP2-induced genomic instability arising predominantly during transcription (Gomez-Herreros et al., 2013; Gomez-Herreros et al., 2017) (Figure 7C,D). In contrast, the observed higher *MLL* breakage and *MLL-AF4* translocations in cells deficient for the MRE11 nuclease activity or for both MRE11 nuclease activity and TDP2, was dependent on both active replication and transcription (Figure 7E,F). MRE11 therefore, appears to suppress *MLL* translocations by preventing accumulation of TOP2-DNA covalent complexes that are converted to DSBs by colliding with replication and transcription machineries. The observed distinct contribution of TDP2 and MRE11 to prevent *MLL* breakage and fusions, was not due to changes in the cell cycle distribution upon the various treatments in these mutants (Figure S7C), but due to predominantly transcription-dependent and replication-independent functions of TDP2 at preventing TOP2-induced DSBs compared to MRE11 (Figure S7D). Taken together, these data indicate that TDP2, the nuclease activity of MRE11 and NHEJ are suppressors of ETO-induced *MLL* translocations by preventing TOP2-induced DSBs or facilitating DSB-repair within the *MLL* gene. In addition to TDP2 activity that averts TOP2-induced DSBs generated during transcription (Gomez-Herreros et al., 2013; Gomez-Herreros et al., 2017), MRE11 nuclease activity prevents the conversion of TOP2-DNA covalent complexes to DSBs both during transcription and replication to suppress *MLL* fusions.

## Discussion

Here, we have systematically investigated how abortive TOP2 functions promote genomic instability and the formation of recurrent leukemia-driving *MLL* rearrangements. We show that transcription and the spatial folding of chromosomes cooperate in the formation of oncogenic *MLL* translocations.

### Transcription, loop anchors and *MLL* translocations

The contribution of transcription to genomic instability mediated by abortive TOP2 activities has been unclear (Canela et al., 2017; Gomez-Herreros et al., 2017; Xiao et al., 2003). Several lines of evidence presented in this study demonstrate that transcription is a major contributor to the occurrence of TOP2-induced DSBs. First, ETO-induced hot spots were highly enriched within active genome regions and genome sites occupied by RNA polymerase II and its active forms. Second, the ETO-induced DNA damage was positively associated with transcriptional output at both promoters and intragenic genome sites. Third, inhibition of transcription elongation led to a substantial decrease on the occurrence of TOP2-induced DSBs across promoters and gene bodies as quantified by both sBLISS and high-throughput imaging.

TOP2 binding (Uuskula-Reimand et al., 2016) and activity (Canela et al., 2017) have been found enriched at boundaries of loop domains. TOP2-induced activity at these sites was shown to be transcription-independent (Canela et al., 2017), and therefore, it was concluded that breaks at these sites may be due to TOP2 enzymes acting to release chromatin entanglements that emerge during loop extrusion dynamics (Canela et al., 2017). We show that loop boundaries coinciding with transcriptionally active genes are vulnerable to TOP2-induced DNA breakage, and that transcriptional activity, the localization at loop anchors and the DNA fragility are strongly interlinked. Likewise, we show that *MLL* and fusion partner genes are highly transcribed, are enriched at loop boundaries, and accumulate TOP2-induced DSBs in transcription-dependent manner. We therefore propose that chromatin loop boundaries are vulnerable to TOP2-induced DSBs, caused by the abortive activities of TOP2 as it dissipates torsional stress arising during transcription. This TOP2-induced DNA fragility at genome sites overlapping with *MLL* and the potential fusion partners’ hot spots drives the formation of oncogenic *MLL* translocations (Figure 8A). Moreover, we observed that DSBs at intragenic CTCF sites localized at loop anchors, are asymmetric, displaying a prominent enrichment between the CTCF sites and the direction of transcription. Stabilized chromatin loops at CTCF/cohesin binding sites at loop anchors represent an obstacle to the dissipation of DNA supercoiling generated during transcription. Therefore, when highly transcribed genes are oriented towards loop anchors, accumulating supercoiling at surmount levels at these regions may arrest transcription. Under these conditions, directed TOP2 activity at the side of the chromatin loop facing toward genes’ TSS (Figure 4E), may cause the observed polarized pattern of chromosome fragility. In line with this view, CTCF binding has been shown to mark boundaries of transcriptionally active supercoiling domains (Naughton et al., 2013) and TOP2 functions at CTCF binding sites to facilitate the remodeling of supercoiling (Uuskula-Reimand et al., 2016). Another possibility is that encounters of the transcription machinery with unresolved TOP2-DNA covalent complexes trigger their conversion to DSBs. In this case, release of covalently linked TOP2 may require the transcription–dependent targeting of TOP2 for degradation as previously shown by biochemical approaches (Ban et al., 2013; Zhang et al., 2006). Alternatively, TOP2 activities at loop anchors might be required to dissipate topological stress associated with the dynamics of loop formation (Canela et al., 2017). However, we did not observe any correlation of the direction of breaks with the direction of loop formation at loop anchors, and the formation of DSBs at these sites was clearly transcription-dependent (Figure 4F). Taken together, we conclude the 3D chromosome organization and transcription are key contributors to DNA fragility at recurrent genome sites that frequently translocate in cancer.

**Figure 8.**
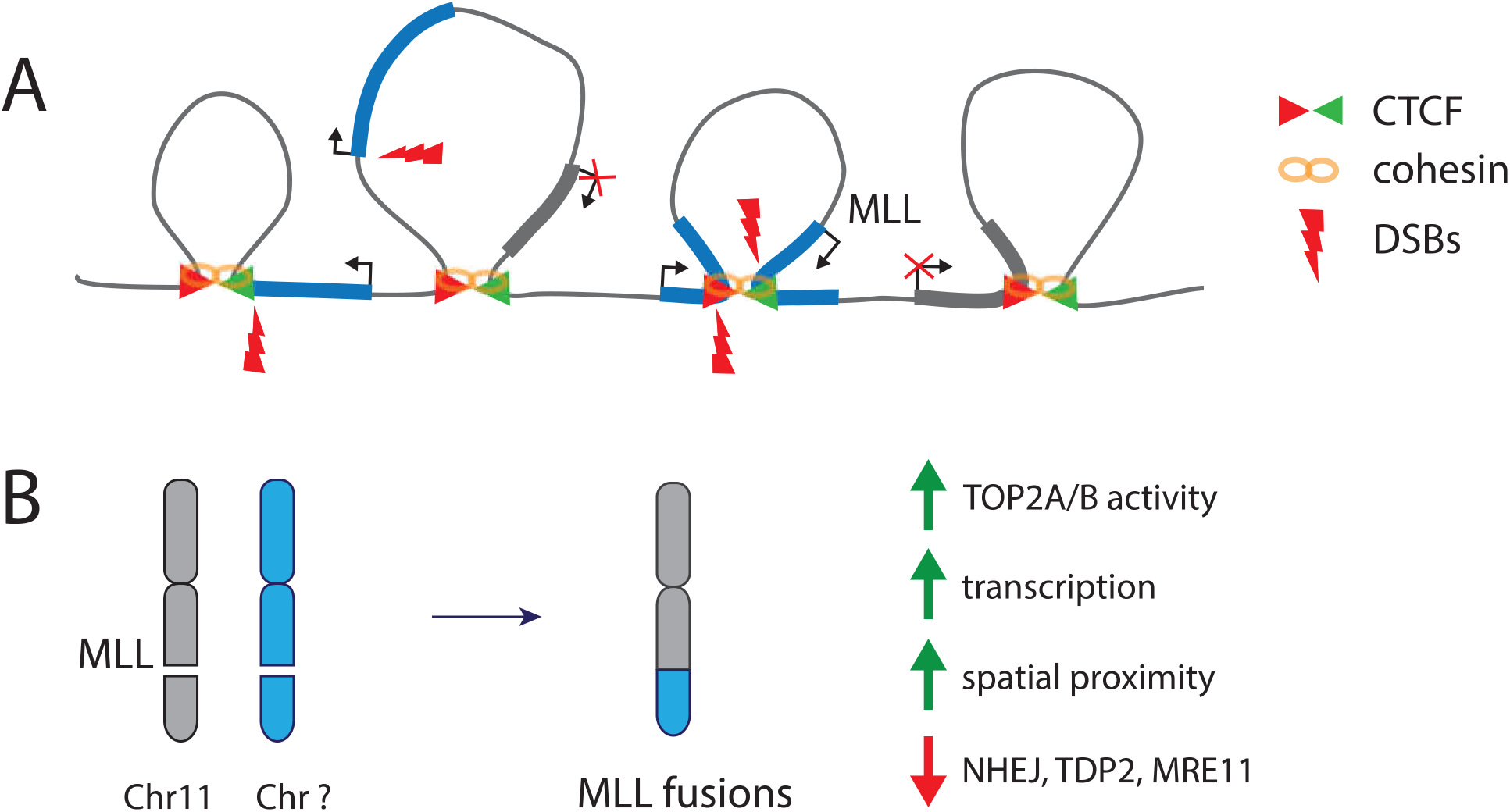
Transcription-driven DNA fragility at loop anchors promotes the formation of *MLL* fusions. (A) TOP2-induced DSBs (red bolts) are enriched at promoters (black arrows) and at CTCF/cohesin sites at loop boundaries (convergent CTCF sites: green/red triangles, cohesin: yellow rings) of active genes (blue lines), but not within inactive genes (grey lines). DSBs at CTCF sites at loop anchors are asymmetric and breakage pattern correlates with the direction of transcription. DSBs within highly transcribed *MLL* and fusion partner genes localized at loop anchor regions, promote the formation of oncogenic *MLL* translocations. (B) ETO-induced DSBs drive chromosomal rearrangements of highly transcribed *MLL* (on chromosome 11, grey) and potential fusion partner genes (various chromosomes, blue) to form oncogenic translocations. The formation of *MLL* fusions is favored by the activities of both TOP2 isoforms, the spatial proximity of *MLL* to partner genes and it depends on transcription elongation. Removal of TOP2-DNA complexes by TDP2 and MRE11 as well as repair of DSBs by NHEJ suppress the formation of oncogenic *MLL* translocations.

### Spatial genome organization and *MLL* translocations

Spatial proximity favors the formation of chromosome translocations (Roukos and Misteli, 2014; Roukos et al., 2013; Zhang et al., 2012), but chromosome damage is a pre-requisite for fusions to occur (Hakim et al., 2012). We show that while common *MLL* fusion partners show similar frequencies of DSBs upon ETO-treatment, *MLL-ENL* fusions arise more often than fusions with other partners, an effect that could be attributed to the closer proximity of *MLL* and *ENL* in hematopoietic cell lines and CD34+ stem cells (Figure 2). However, while *MLL-ENL* fusions occur more often, the predominant fusion found in patients with t-AML is *MLL-AF9* (Meyer et al., 2018), suggesting that the selective advantage conferred by certain fusions during oncogenesis, rather than their formation frequencies, govern their prevalence in patients with leukemias. In accordance, the various *MLL* fusions have different transforming capacities *in vivo* and these are closely linked to microenvironment (Drynan et al., 2005; Krivtsov and Armstrong, 2007; Lin et al., 2017).

Previous studies have implicated transcription as a facilitator of *MLL* fusion partner synapsis (Cowell et al., 2012). Our analysis of the spatial arrangement of the potential fusion partners in single cells showed that transcription does not contribute to the spatial proximity of fusion partners in absence of DNA damage. Instead, we observed that transcription-dependent TOP2-induced chromosome breakage could further promote the synapsis of chromosome breaks, only between proximal partners in three-dimensional nuclear space. These findings indicate that *MLL* fusions occur predominantly between genes pre-synapsed in the three-dimensional nuclear space and that break mobility may further promote gene synapsis and translocations, only between already spatially proximal genes, such as *MLL* and *ENL.*

### Contribution of TOP2-isoforms to the formation of DSBs and *MLL* fusions

While topoisomerase poisons can equally trap both TOP2 isoforms (Cornarotti et al., 1996; Nitiss, 2009; Willmore et al., 1998), the genomic instability associated with TOP2-targeted drugs has been attributed predominantly to the TOP2B isoform (Azarova et al., 2010; Azarova et al., 2007; Canela et al., 2017; Cowell et al., 2012). However, different studies have demonstrated TOP2A isoform-dependent functions (de Campos-Nebel et al., 2010; Tammaro et al., 2013) and trapped-TOP2A cleavage complexes have been detected within *MLL* translocation hot spots and of frequently translocating genome loci (Yu et al., 2017). Our systematic analysis upon loss-of-function experiments in different cell lines and the t-AML relevant CD34+ stem cells, show that, although to varying degrees, both TOP2A and TOP2B isoforms contribute to ETO-induced DNA damage and the formation of *MLL* fusions (Figure 8B). The observed variability in contribution can be accredited to the different relative levels of the TOP2-isoforms in the various cell lines, clarifying previous findings (Azarova et al., 2010; Azarova et al., 2007; Canela et al., 2017; Cowell et al., 2012; de Campos-Nebel et al., 2010; Tammaro et al., 2013). In support of this idea, ETO-induced DNA damage, *MLL* breakage and *MLL* fusions (Figure 6F,G), occur more often in S and G2 phases of the cell cycle, when both TOP2 isoforms are expressed.

### Suppressors of *MLL* fusions

We have systematically addressed the contribution of factors involved in removal of TOP2ccs and of the subsequent DSB-repair pathways on the susceptibility of the *MLL* locus to breakage and the formation of ETO-induced *MLL* translocations. Cells defective for HR show no apparent differences in the fraction of cells with ETO-induced breaks within the *MLL* locus and with *MLL* fusions, findings consistent with the current notion that NHEJ is the major pathway repairing TOP2-induced DSBs (Maede et al., 2014; Malik et al., 2006). In line, deficiency of NHEJ led to an increase of ETO-induced breaks within the *MLL* locus, which promoted the formation of *MLL* fusions (Figure 7A,B). These findings suggest that NHEJ acts as a suppressor of oncogenic *MLL* fusions, by mediating rapid intrachromosomal repair of TOP2-induced breaks within frequently translocating genes. In its absence, persistent unrepaired TOP2-induced DSBs may have a longer time window to pair with other DSBs and to translocate, while the final ligation step can be mediated by the alternative-NHEJ (alt-NHEJ) pathway utilizing the activities of other ligases such as LIG3 and/ or LIG1 (Simsek et al., 2011; Soni et al., 2014). These observations have important implications for cancer therapy, since inhibition of NHEJ has been proposed to potentiate TOP2 poison-induced cytotoxicity (Willmore et al., 2004; Zhao et al., 2006). Although inhibiting NHEJ would synergize with TOP2 poison treatment to effectively kill cancer cells, it may also increase the occurrence of oncogenic fusions that lead to secondary, therapy-related cancers.

We furthermore addressed the impact of factors that are involved in removal of TOP2ccs on *MLL* susceptibility to breakage and translocations. In agreement with previous studies (Gomez-Herreros et al., 2017), TDP2 prevented the accumulation of ETO-induced DSBs within the *MLL* locus and the formation of *MLL* fusions (Figure 7A,B), an effect that can be attributed to TDP2 functions in removing TOP2ccs accumulating predominantly during transcription (Figure 7C,D). Interestingly, deficiency of TDP2 did not lead to genomic instability in absence to ETO-treatment, suggesting that endogenous TOP2ccs can be removed by additional alternative mechanisms.

Indeed, we found that the nuclease activity of MRE11 suppresses TOP2-induced DSBs within *MLL* and the formation of subsequent *MLL* fusions, by acting in a separate to the TDP2 pathway (Figure 7A,B). Importantly, in contrast to the transcription-specific functions of TDP2 in preventing breakage within *MLL*, we found that the translocation events that are suppressed by nuclease activity of MRE11, were dependent on both replication and transcription (Figure 7E,F). This finding suggests that MRE11 is able to remove TOP2ccs from DNA by acting during replication, in addition to G1 phase (Hoa et al., 2016; Quennet et al., 2011). In summary, TDP2, the nuclease activity of MRE11 and the NHEJ pathway, suppress oncogenic *MLL* translocations, by either preventing the conversion of TOP2ccs to DSBs within *MLL* and potential translocation partners or by facilitating intrachromosomal DSB-repair, respectively (Figure 8B).

Taken together, we conclude that an intriguing interplay, between transcriptional activity and spatial genome organization, determine the occurrence of *MLL* chromosomal translocations. Our findings provide a mechanism by which transcription-associated changes of DNA topology at loop boundaries contribute to genomic instability driving oncogenic translocations, and uncover key molecular players that actively suppress them.

## Author contributions

The study was conceived and designed by V.R. with input from H.J.G., E.G.G. and A.P. H.J.G. performed all experiments except sBLISS (B.A.M.B.), the experiments assessing the contribution of replication/transcription to ETO-induced DSBs by imaging and the experiments assessing the role of TOP2 isoforms on ETO-induced DSBs (R.P. and V.M). Bioinformatics analyses performed by E.G.G. with the help of S.S. and G.P, under the supervision of A.P. and V.R. Code for C-Fusion 3D analysis was generated by O.D., and N.J. and A.M. performed the transcription factory RNA-seq experiments and analysis. C.F.N. and D.F.H. generated the HCT116 inducible TOP2A knockout cell line and S.T. and H.S. the various repair mutants in TK6 cells. E.M.W. and T.K. isolated CD34+ cells. L.B. contributed to interpretation of data. The manuscript was written by V.R. with input from all authors.

## Acknowledgements

We thank Reza Mirzazadeh (N.C. lab) for initial help with BLISS, Silvano Garnerone (N.C. lab) for support with processing raw BLISS data, and Prof. Andy Porter and Leszek Wojnowski for sharing reagents. Work in V.R. lab is supported by the Naturwissenschaftlich-medizinisches Forschungszentrum (NMFZ) grant and in N.C lab by the Swedish Research Council (521-2014-2866), the Swedish Cancer Research Foundation (CAN 2015/585), the Ragnar Söderberg Foundation, and the Strategic Research Programme in Cancer (StratCan) at Karolinska Institutet. Work in A.P. lab is supported by CMMC core funding and by the German Ministry for Research (DFG; grant no. PA2456/4-1); in H.S. lab by the Takeda research and Mitsubishi foundation; in S.T. lab by the JSPS Core-to-Core Program, A. Advanced Research Networks, and in the D.F.H. lab by National Health and Medical Research Council (Australia) project (Grants GNT1127209 and GNT1145188) and by the Victorian Government’s Operational Infrastructure Support Program. B.B. was supported by a Rubicon postdoctoral scholarship from NOW in the Netherlands. The Opera Phenix High Content Screening System is supported by the “DFG Major Research Instrumentation Programme” (INST 247/845-1 FUGG). Support by the Microscopy and Histology Core Facility and the Genomics Core Facility and the use of its NextSeq500 (INST 247/870-1 FUGG) is also gratefully acknowledged.

## Declaration of interests

The authors have no competing interests to declare.

## Materials and correspondence

Requests for materials should be addressed to Vassilis Roukos

## Data availability

All sequencing data have been submitted to the Gene Expression Omnibus under the accession number GSE121742.

## Supplementary Figure legends

**Figure S1.**
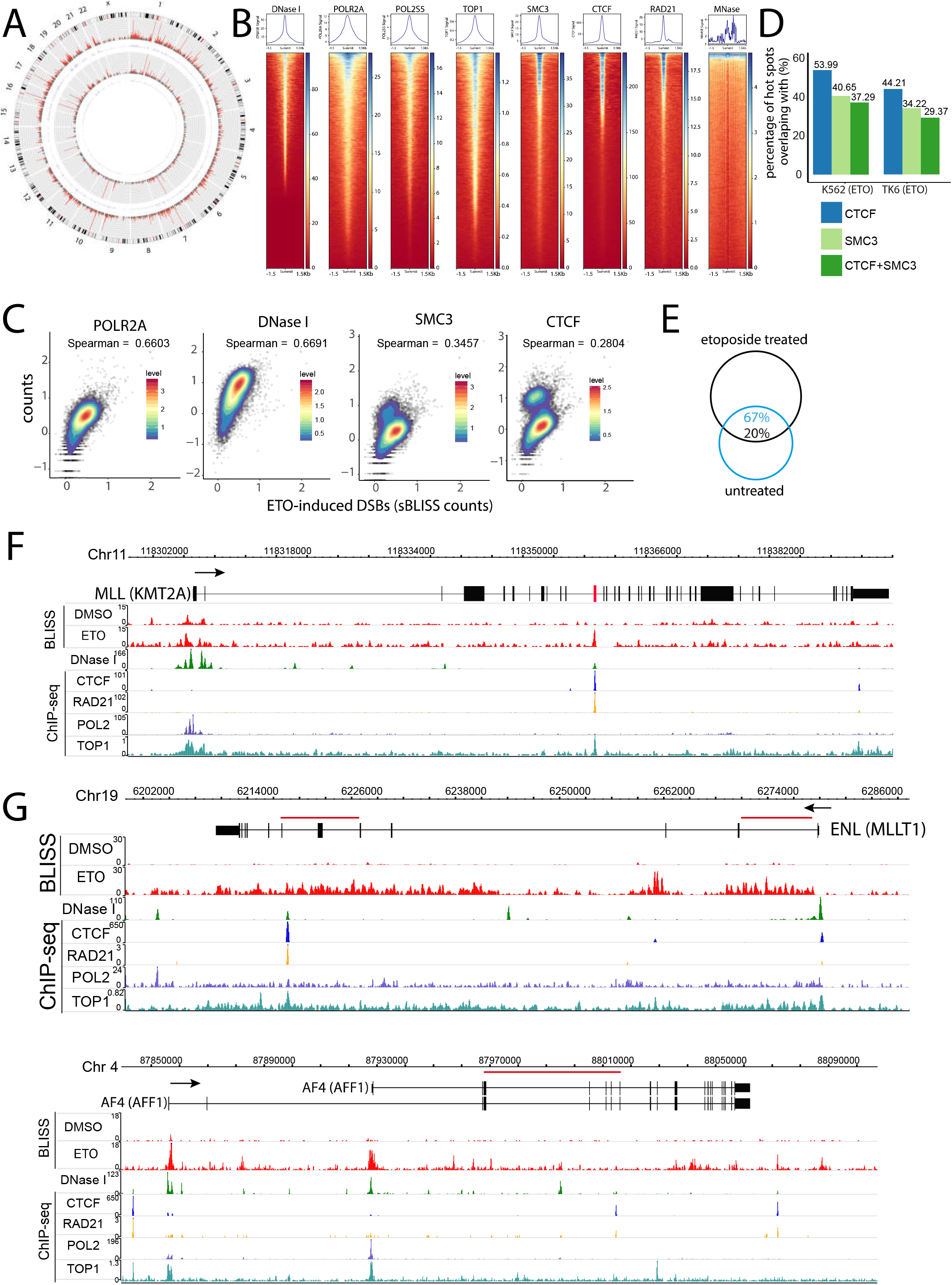
Profiling ETO-induced DSBs across the genome. (A) Circos plot depicting DSB hotspots detected by sBLISS in DMSO-(inner two circles) or ETO-treated (20μM, 6h) K562 cells (two outer circles). Grey tracks mark hot spots with adjusted p<0.05, red tracks with adjusted p<1e-10. Breakage hotspots overlapping with centromeric regions were excluded from visualization. (B) Aggregate plots and signal heatmaps of indicated features centered on ETO-induced DSB hotspots in K562 cells, extended for a total of 3Kbp from its summit. (C) Spearman’s correlation coefficient between ETO-induced DSBs and ChIP-seq or DNase I-seq read counts (RPKM) of indicated features per gene in K562 cells. Color key represents a two-dimensional kernel density estimation. (D) Percentage of ETO-induced DSB hot spots overlapping with ChIP-seq peak signals of the indicated proteins. (E) Venn diagram showing the overlap in percentage of spontaneous and ETO-induced hot spots in K562 cells. (F) sBLISS DSB-profiles along the *MLL (KMT2A)* gene in DMSO and ETO-treated (20μM, 6h) K562 cells. Exons are shown as black squares, the t-AML translocation hot spot in *MLL* is marked red (G) DSB-mapping by sBLISS within *ENL (MLLT1)* and *AF4 (AFF1)* genes in DMSO or ETO-treated (30μM, 4h) TK6 cells. Red lines correspond to BCRs in these genes. ChIP-seq data were derived from ENCODE (Davis et al., 2018) and (Baranello et al., 2016). Arrows represent the direction of transcription.

**Figure S2.**
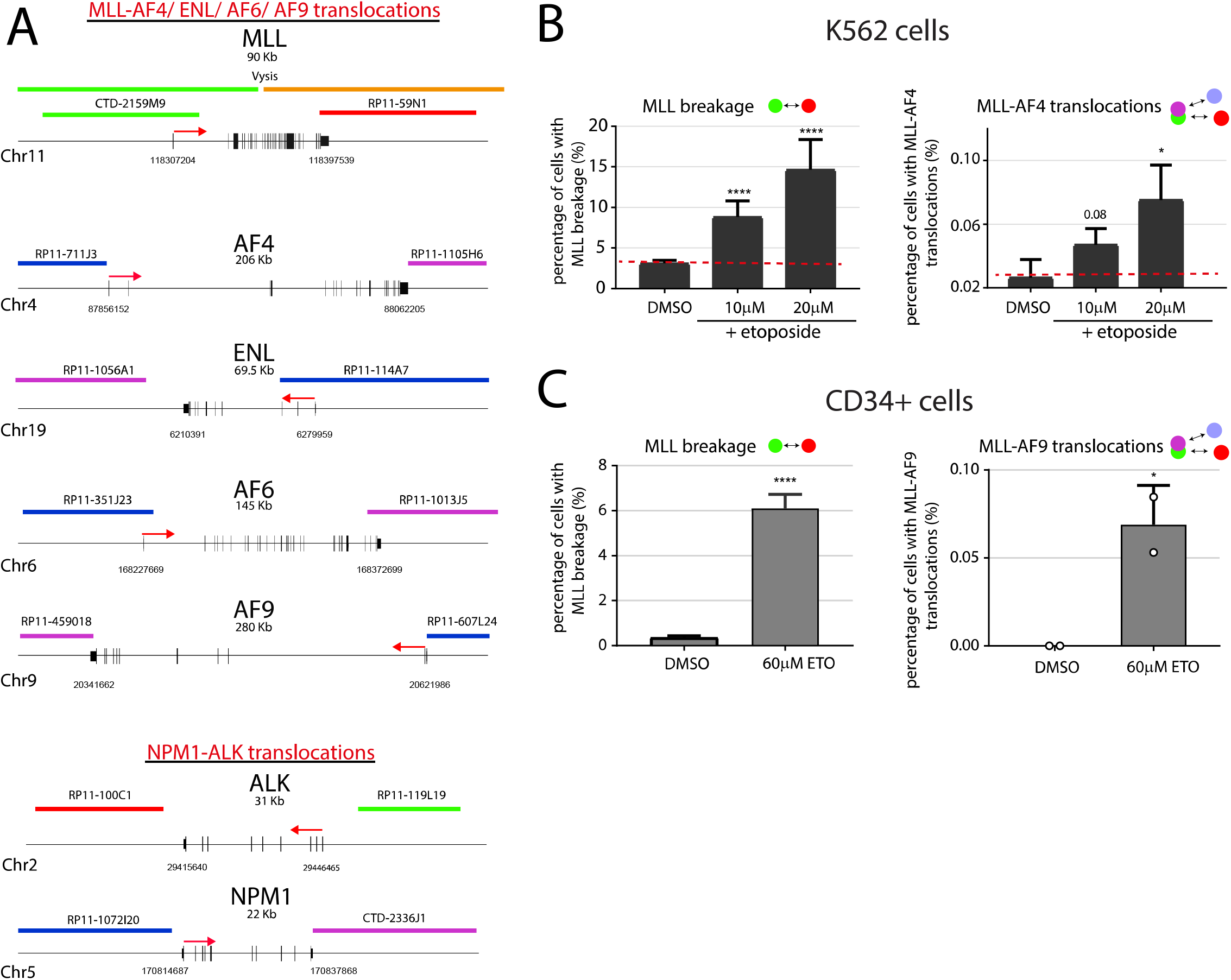
ETO-treatment induces the formation of *MLL* translocations in human hematopoietic cell lines and CD34+ progenitor cells. (A) Schematic representation of genes and FISH probes used in this study. (B) *MLL* breakage and *MLL-AF4* translocation frequencies in K562 cells, and (C) *MLL* breakage and *MLL-AF9* translocation frequencies in CD34+ progenitor cells measured by 4-color C-Fusion 3D upon release for two days after ETO-treatment at the indicated doses for 4h.

**Figure S3.**
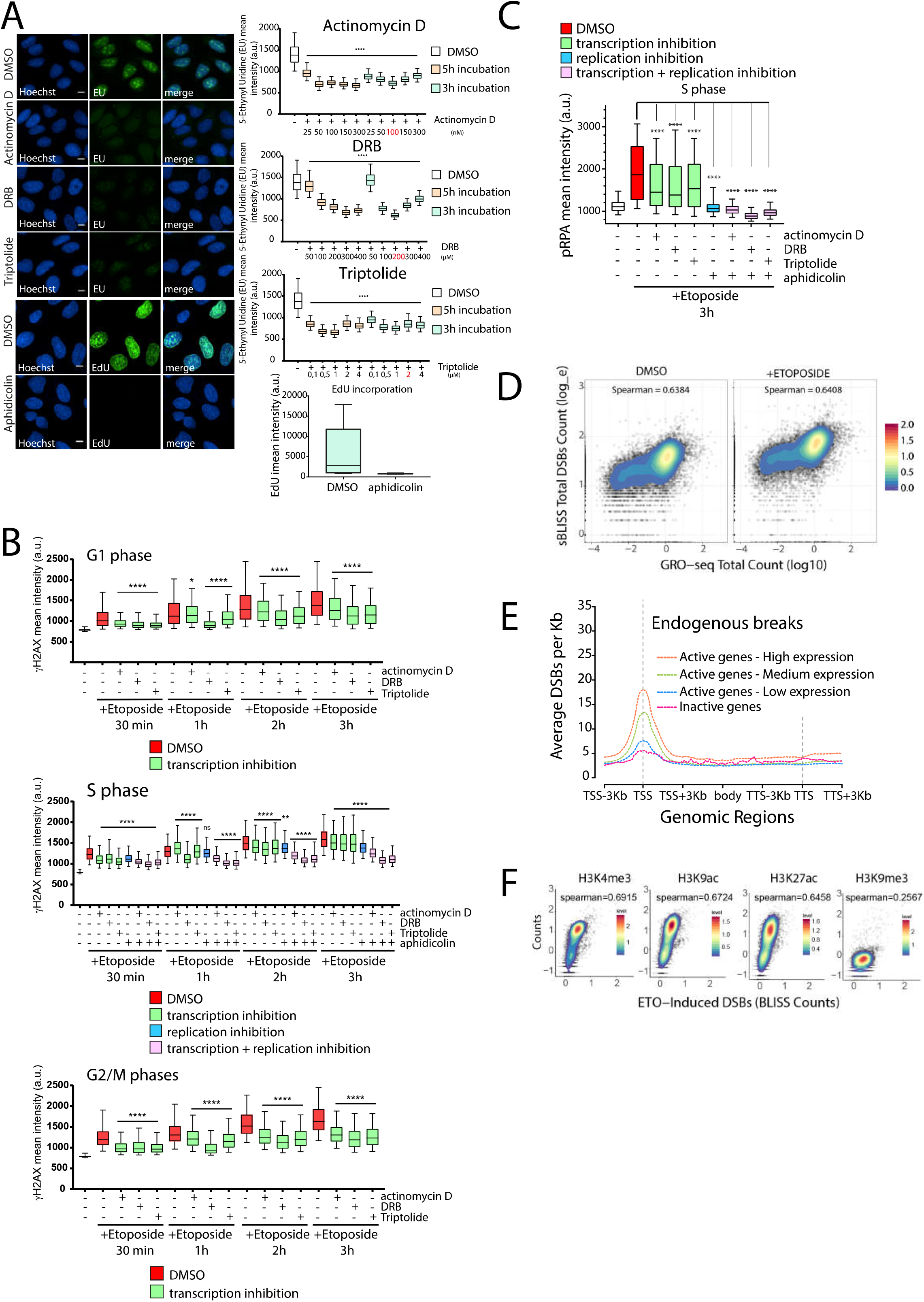
Transcription contributes to ETO-induced DNA breakage. (A) U2OS cells were pretreated with the indicated doses of actinomycin, DRB or triptolide for 3 or 5 hours, and nascent RNA expression was assessed by quantification of 5-Ethynyl Uridine (EU) incorporation. Similarly, inhibition of replication was evaluated by assessing levels of active replication by EdU incorporation upon treatment with 20μM aphidicolin for 3h hours. Selected drug dosages for further experiments are marked red. Image scale bars correspond to 10μm. (B) Contribution of transcription and/or replication on ETO-induced DSBs across the cell cycle. Experiment performed as in Figure 3A. The cell cycle staging of individual cells was assessed by quantifying the DNA content by Hoechst staining as previously shown (Roukos et al., 2015). (C) Inhibition of transcription (pretreatment for 3h with actinomycin D: 100nM, DRB: 200μM, triptolide: 2μM, and/ or replication, aphidicolin: 20μM) in S phase of U2OS cells led to a decrease in ETO-induced (10μM for 3h) immunostaining of phospho-RPA (phospho-Ser4/Ser8), a marker of collapsed replication forks (Toledo et al., 2013) (at least 1000 cells were analyzed in each sample, ****P < 0.0001, One Way ANOVA test and Tukey test). (D) Spearman’s correlation coefficient between DSB counts measured by sBLISS of DMSO treated (endogenous DSBs, left) or ETO-treated (20μM, 6h, right) K562 cells and the total GRO-seq read count (RPKM) per gene. The color key represents a two-dimensional kernel density estimation. (E) Aggregate plot of average DSBs per Kbp in DMSO-treated K562 cells (endogenous DSBs) across genes classified by expression as in Figure 3C. (F) Spearman’s correlation coefficient between DSB-counts and the indicated histone modification ChIP-seq read counts (RPKM) per ETO-induced hotspot in K562 cells (20μM, 6h). The color key represents a two-dimensional kernel density estimation.

**Figure S4.**
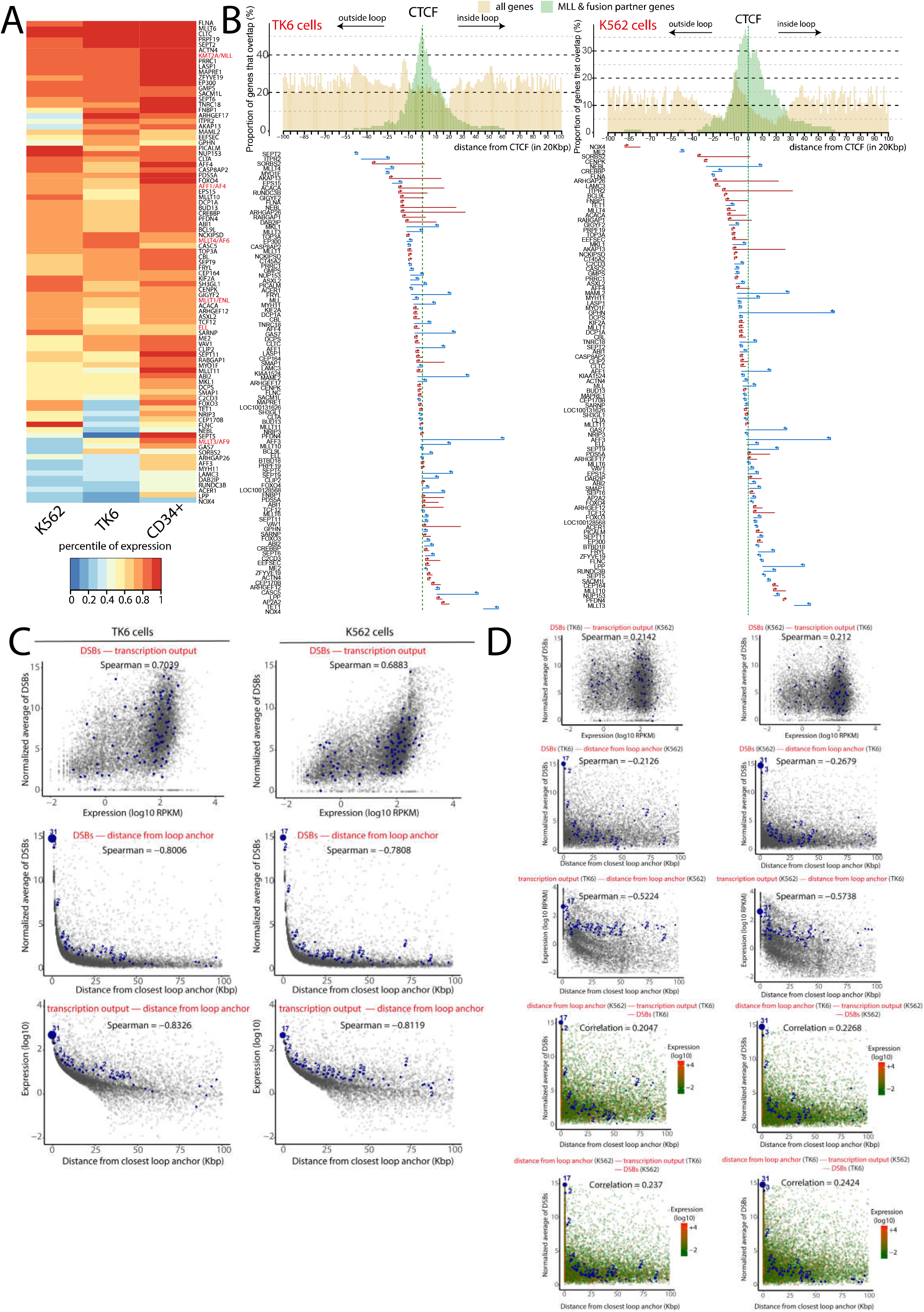
*MLL* and fusion partner genes are localized at loop anchors, are highly transcribed and accumulate high levels of TOP2-induced DSBs. (A) Heatmap displaying expression values of the *MLL* and fusion partners genes (Meyer et al., 2018) given as percentiles of the genome wide expression levels in hematopoietic K562 (GRO-seq) and TK6 cells (nascent RNA (Melnik et al., 2016)) and in progenitor CD34+ cells (RNA-seq). (B) Histogram of the proportion of overlapping gene bodies of *MLL* and fusion partners genes (green) or control genes (all genes in the genome; yellow) relative to loop anchor sites marked by CTCF, in TK6 and K562 cells. Genes were placed with regard to their respective distance between the closest loop anchor’s CTCF position and the gene’s TSS. Bottom panel shows a visual depiction of the orientation of *MLL* and fusion partners’ genes around the centered CTCF position. Arrows on gene bodies show transcription directionality. Genes in which the closest loop anchor was the downstream anchor of a chromatin loop were mirrored and are depicted in their opposite orientation. (C) Correlations between transcription output, distance from their closest loop anchor and average ETO-induced DSB counts per gene for TK6 and K562 cells. Distances were rounded up to the nearest Kbp and were measured from the closest gene end (TSS or TTS) to the middle of the closest loop anchor; distances of 0 bp correspond to an overlap between the gene and the 10-Kbp loop anchor. *MLL* and fusion partner genes are represented as blue circles and the size corresponds to the number of overlapping genes (number is shown on top). (D) Cross cell line correlations between transcription output, distance from their closest loop anchor and average ETO-induced DSB counts per gene. Data on these features derived from K562 and TK6 cells were exchanged as indicated. *MLL* and fusion partner genes are indicated by blue circles as in (C).

**Figure S5.**
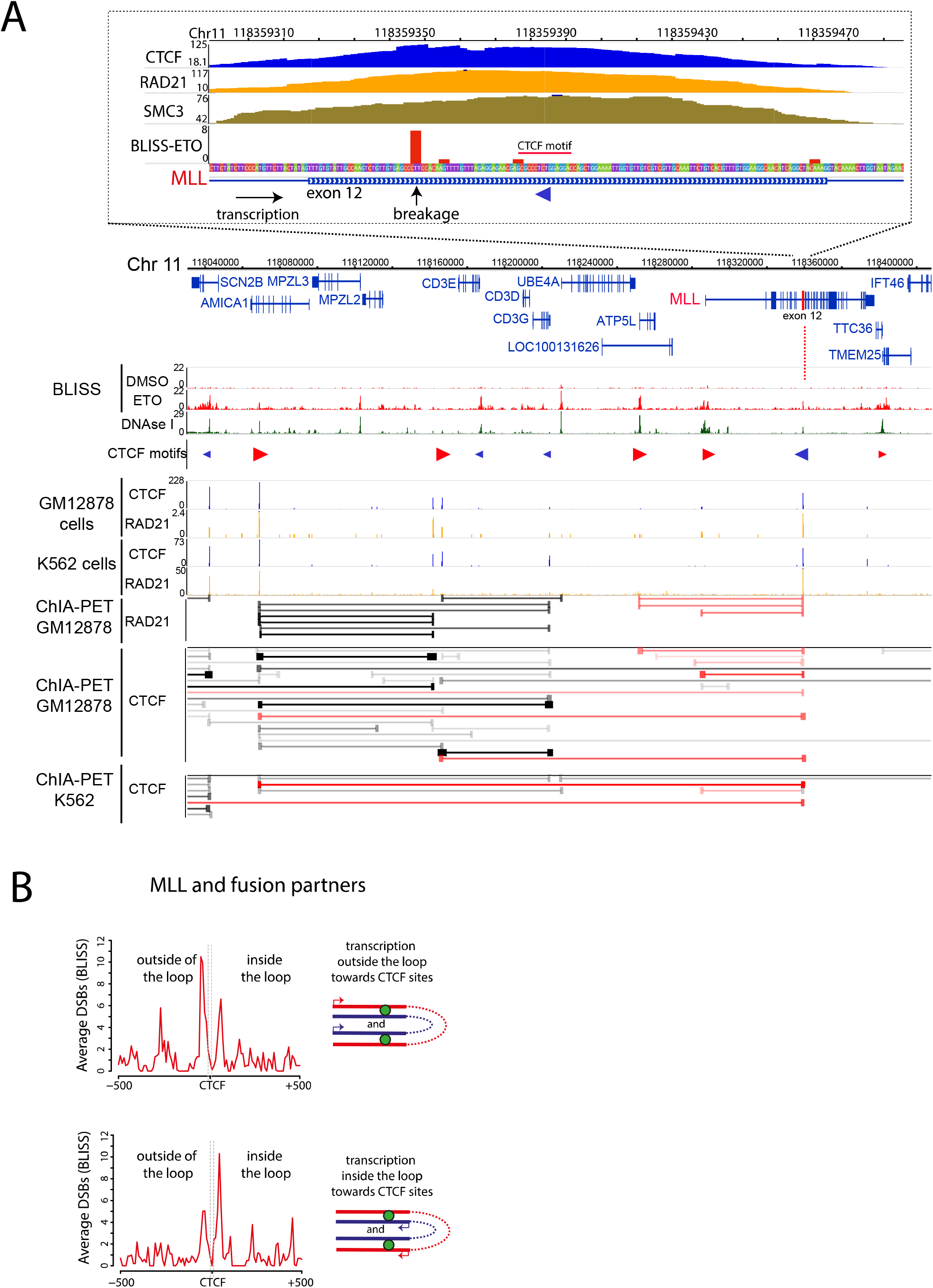
DSB-patterns at CTCF sites within *MLL* and fusion partner genes located at loop anchors correlate with transcription directionality. (A) The *MLL* hot spot is localized at the border of a chromatin loop and the DSBs are localized within the loop, between the CTCF site and the direction of transcription. DSB profiles upon ETO-treatment around the human *MLL* locus (TK6 cells). CTCF motifs (G-rich, forward strand=red, reverse strand=blue) and CTCF and cohesin subunit Rad21 binding sites (ChIP-seq) are shown. Chromatin loop interactions measured by CTCF, RAD21 ChIA-PET are indicated by horizontal lines (pink lines indicate interaction emanating from the *MLL* hot spot with upstream genomic loci). Inset above shows in magnification the localization of DSBs at the *MLL* translocation hot spot. Note that the breakage is within the chromatin loop (between the CTCF motif (red triangle) and the direction of transcription (black arrow). (B) As in Figure 4E, only for *MLL* and fusion partner genes located within loop anchor sites in ETO-treated K562 cells.

**Figure S6.**
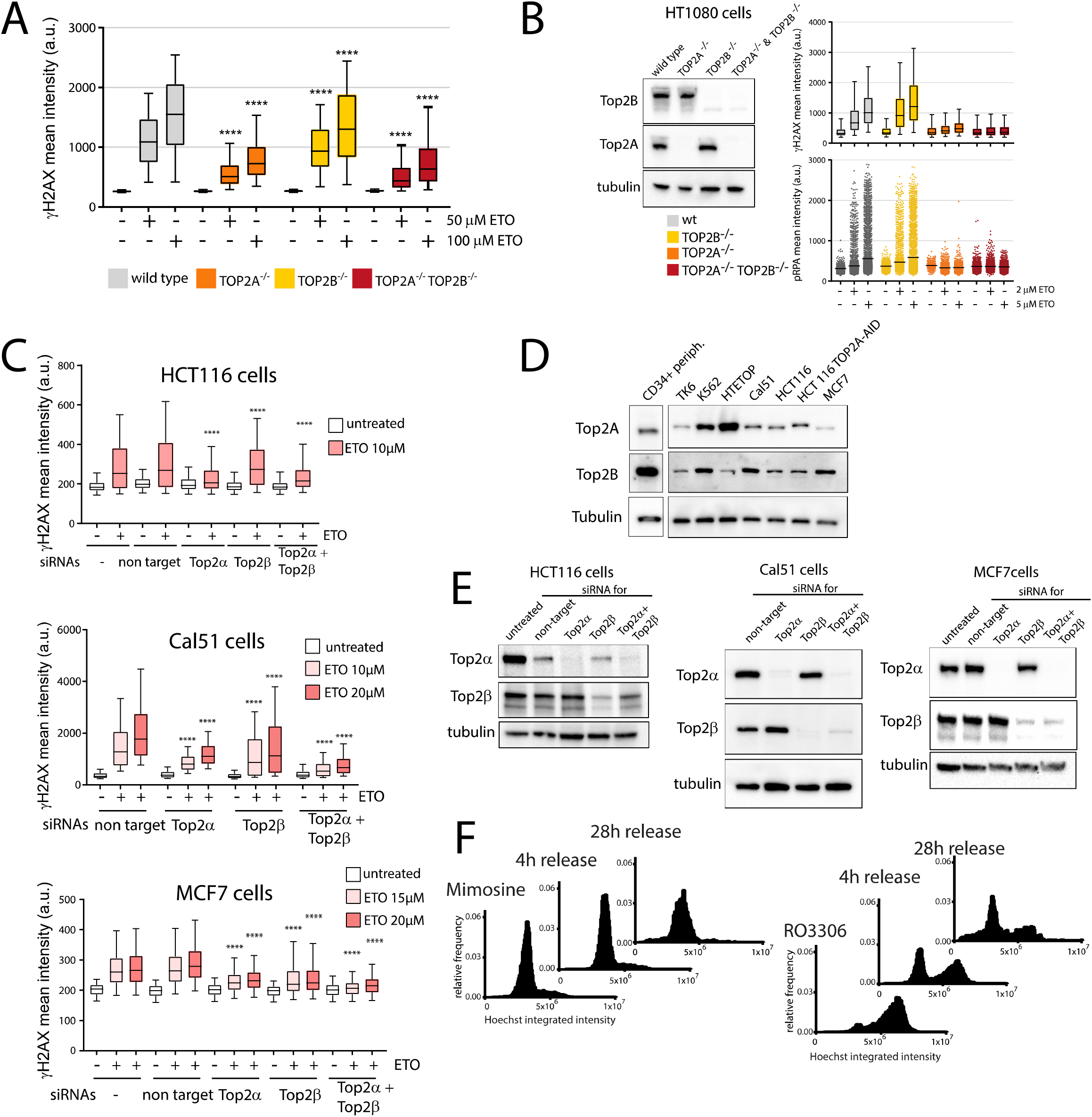
Contribution of TOP2-isoforms to ETO-induced DSBs. (A) Quantification of ETO-induced DSB-signaling in HCT116 cells by γH2AX immunostaining in the absence of TOP2 isoforms (4500 to 8400 cells were analyzed per sample, one representative out of three independent experiments is shown. *P < 0.05, **P < 0.01, ***P < 0.001, ****P < 0.0001, One Way ANOVA and Tukey test comparison to wildtype cells). (B) TOP2A and/or TOP2B expression levels in HT1080 fibrosarcoma cells (HTETOP) deficient for TOP2A and/or TOP2B by western blotting. Quantification of γH2AX or pRPA (phopho-Ser4/Se8) immunostaining levels upon etoposide are shown in the absence of the respective TOP2 isoforms. **(C)** ETO-induced breakage was assessed by quantifying γH2AX phosphorylation levels in the indicated cell lines upon siRNA-mediated knock down of TOP2A and/ or TOP2B expression. Time and concentration of ETO-treatment was optimized for each cell line (Cal51 cells: etoposide 10μM and 20μM for 4h, MCF-7 cells: etoposide 15μM and 20μM for 3h). Statistical significance was measured by One Way ANOVA test and Tukey test. Quantification of the number of γH2AX foci led to similar results (data not shown). (D) Expression levels of TOP2A and TOP2B in various human cell lines and peripheral hematopoietic CD34+ stem cells and progenitors. (E) The efficiency of knock down was evaluated by western blotting 3days after siRNAs transfection. (F) Cell cycle distributions of cells released from G1 (Mimosine) or G2/M (RO3306) in the presence of 20μM ETO (4h release) and after ETO treatment for an additional 24h (28h release) in the presence of the second inhibitor.

**Figure S7.**
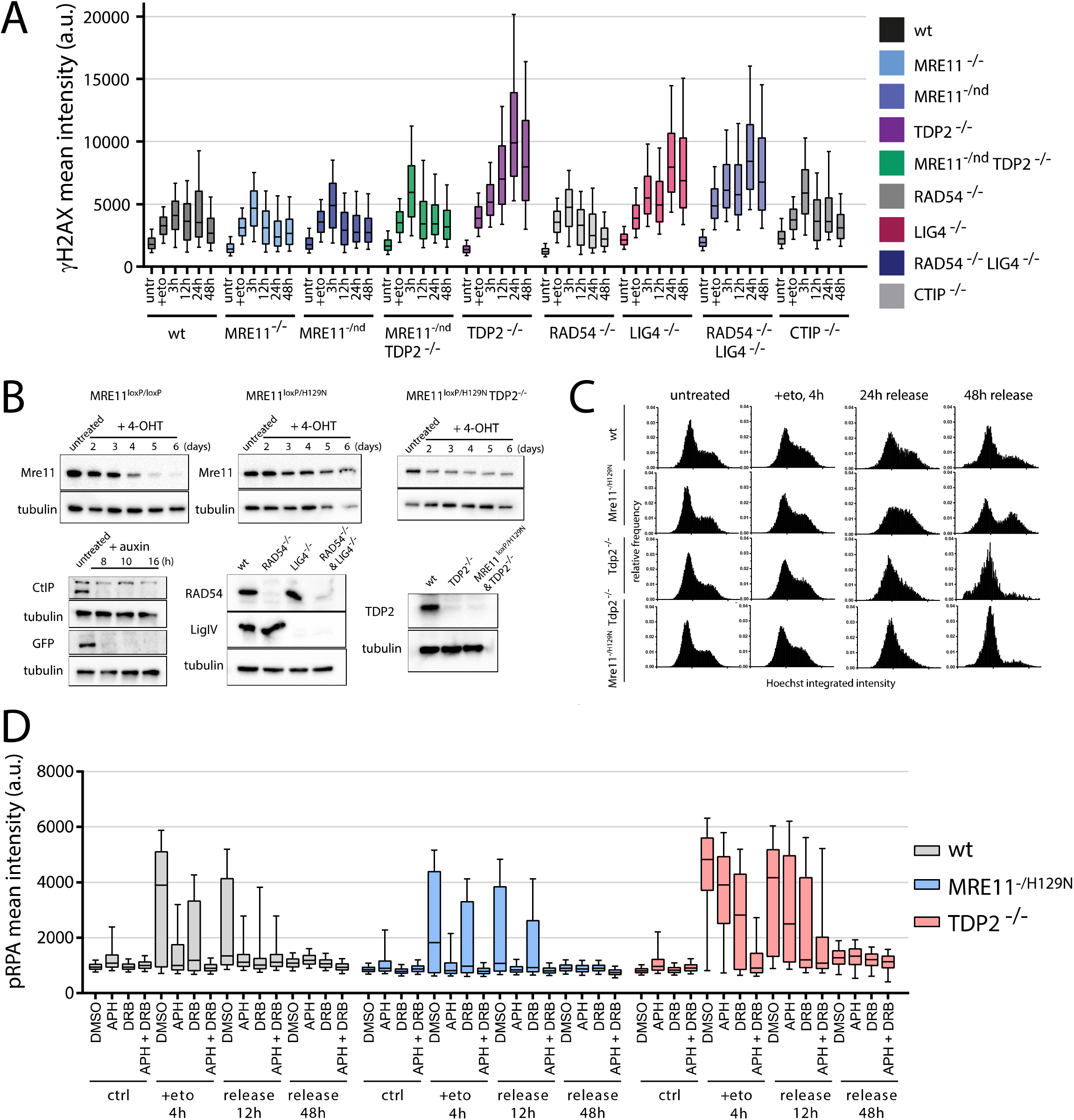
Characterization of repair kinetics in cells deficient for factors involved in the removal of TOP2-DNA covalent complexes and DSB repair. (A) The indicated deficient TK6 cell lines were treated with 20μM ETO for 4h and released for 3h up to two days. Repair kinetics in the different mutant cell lines were assessed by quantifying levels of γH2AX phosphorylation. Note that deficiency in MRE11 nuclease activity results to reduced γH2AX phosphorylation upon ETO-treatment even in the MRE11^−/H129N^TDP2^−/-^ double mutant compared to TDP2^−/-^, which may indicate inhibition of signaling towards H2AX (750 to 35000 cells were analyzed per sample). (B) Validation of the constitutive or inducible knock-out cell lines used in experiments described in Figure 7 and S7 by western blotting. (C) Cell cycle profiles of the indicated mutants upon treatment and release for up to 2 days from ETO treatment (10μM, 4h). (D) The indicated deficient cell lines were treated as in Figures 7C-E and immunostaining for pRPA (phospho-Ser4/Ser8) was quantified by high-throughput imaging and automated image analysis (2000 to 12,500 cells were analyzed per sample).

## Methods

### Generation of knockout cell lines

The HTETOP cell line (derived from the fibrosarcoma human cell line HT1080) has been previously described (Carpenter and Porter, 2004). Mre11^loxP/loxP^, Mre11^loxP/H129N^, TDP2^−^/^−^, Mre11^loxP/H129N^ TDP2^−/-^, LigIV^−/-^, RAD54^−/-^, LigIV^−/-^ RAD54^−/-^ and CtIP auxin-degron lymphoblastoid TK6 cells have been previously characterized (Hoa et al., 2015; Hoa et al., 2016). For the generation of the HCT116-TOP2A-mAID auxin-inducible degron cells, guide RNAs targeting the 3’ UTR of the TOP2A gene were designed using http://crispr.mit.edu/ (Zhang Lab, MIT 2013) with Bbs1/Bpi1 overhangs and ligated into pSpCas9(BB)-2A-GFP (PX458) (#48138, Addgene), a gift from Feng Zhang. Homology directed repair templates were assembled using NEBuilder HiFi DNA Assembly Master Mix (NEB): 300bp homology arms flanking the TOP2A 3’UTR overwriting the stop codon, either mAID-HYGRO or mAID-BSR pieces cut by BamHI from plasmids pMK287 and pMK288 (gifts from Masato Kanemaki NIG Japan, #72825 and #72826, Addgene), and pSK_BS+ cut by EcorI as backbone. Plasmids were transfected into HCT116 +CMV-OsTIR1 cells (a gift from Masato Kanemaki) using Lipofectamine 3000 (Thermo-Fisher) according to manufacturer protocol. 24 h after transfection, cells were reseeded and grown with 125ug/ mL Hygromycin and 7.5ug/ mL Blasticidine for >12 days before colony selection. Correct clones were confirmed by PCR, immunoblotting and immunofluorescence microscopy (TOP2A antibody, sc-166934 Santa Cruz).

HTETOP knockout cells for TOP2B were generated by co-infection of lentiviral particles generated by transfection in HEK293T cells of plasmids expressing SpCas9 (plentiCRISPR v2 (Addgene #52961) and two guide RNAs targeting the TOP2B exon 1 (5’-CGCGCCGCAGCCACCCGACT-3’) and exon 2 (5’-CTTCGTCCTGATACATATAT-3’). For HCTll6-TOP2A-mAID cells, pL-CRISPR.EFS.GFP (Addgene #57818) expressing SpCas9 and pMuLE Lenti Dest Neo (Addgene #62178) containing both of the above TOP2B guide RNAs were used. For the generation of lentiviral particles, HEK293T cells were co-transfected with the second generation packaging vectors (psPax2, pMD2.G) and the lentiviral vector of interest. The lentiviral containing supernatant was filtered and concentrated 10-100X (Lenti-X™ Concentrator, Clontech). Cells were transduced with concentrated virus and 5μg/ mL polybrene (TR-1003-G, Merck) at 800g for 30 min. The following day, transduced cells were enriched with the following antibiotics for at least 10 days: 0.5μg/ mL puromycin for HTETOP cells, 700μg/ mL G418 for HCT116-TOP2A-mAID cells. Single clones were obtained and screened for the knockout of TOP2B by immunofluorescence using the polyclonal TOP2B antibody H-286 (Santa Cruz). They were further characterized by immunoblotting, using the TOP2B antibodies H-286 and A-12 (Santa Cruz).

### Maintenance of cell lines

TK6 cells were cultured in RPMI (Gibco 11875093) supplemented with 5% horse serum (Gibco 11510516), 1 mM sodium pyruvate (Gibco 11360070),100 U/mL penicillin-streptomycin (Gibco 15140122) and 2mM L-glutamine (Gibco 25030081) at 37 °C under a humidified atmosphere with 5% CO_2_. K562 and HCT116 cells were maintained in RPMI supplemented with 10% FBS (Gibco 10270106), 100 U/mL penicillin/ streptomycin and 2mM L-glutamine. HTETOP cells were grown in RPMI supplemented with 10% FBS, 100 U/mL penicillin/ streptomycin and 2mM L-glutamine, 20mM HEPES (Gibco 15630080) and 4x MEM non-essential amino acids (Gibco 11140035). MCF-7, HEK293T, Cal51 and U2OS cells were cultured in DMEM (Gibco 41965062) supplemented with 10% FBS, 100 U/mL penicillin/ streptomycin and 2mM L-glutamine.

Bone marrow CD34+ hematopoietic stem and progenitor cells were purchased from Lonza and maintained in StemSpan SFEM II (Stem Cell Technologies 09605) supplemented with SCF 100ng/ mL, FLT3 50ng/ mL, TPO 50ng/ mL, IL3 20ng/ mL (Peprotech).

To obtain peripheral CD34+ cells, heparin-treated peripheral blood (PB) samples were obtained from healthy donors stimulated with granulocyte-colony stimulating factor (G-CSF) or patients in complete remission proceeding to autologous stem cell transplantation for their primary disease at the University Medical Center of Mainz, Germany. Informed consent was obtained in accordance with the Declaration of Helsinki, and laboratory experiments were performed with approval from the local ethics committee institutional review boards [837.270.05 (4928)]. Mononuclear cells (MNCs) were isolated by means of Ficoll-Hypaque (Seromed) density-gradient centrifugation, re-suspended in freezing media (90% fetal bovine serum (FBS; Biochrom GmbH), 10% dimethylsulfoxide (DMSO; Sigma) and stored in liquid nitrogen. To isolate CD34+ hematopoietic progenitor cells, frozen samples were thawed, washed twice and maintained in IMDM media (Sigma-Aldrich) on ice. Dead cells were eliminated using the Dead Cell Removal Kit (Miltenyi Biotec) and CD34+ cells were enriched by positive selection using the CD34 MicroBead Kit (Miltenyi Biotec) according to the manufacturer’s protocol. Purity of isolated cells was evaluated by flow cytometry upon labeling with an anti-CD34-PE antibody (Miltenyi Biotec) and samples with a purity of > 95% were used for further experiments.

In HTETOP cells the conditional knockout of TOP2A was induced by 10μg/ mL doxycycline (D9891, Sigma-Aldrich) for 48h prior to experiments. HCT116-TOP2A-mAID and HCT116-TOP2A-mAID-TOP2B^−/-^ were maintained in 1μg/ mL puromycin, 125μg/ mL hygromycin and 7.5μg/ mL blasticidin (Invivogen) and TOP2A was triggered to be degraded by addition of 500μM auxin (I5148, Sigma-Aldrich) for 4h. CtIP depletion was obtained by treatment of TK6 degron cells with 1mM auxin for 6h prior to experiments.

### Drug treatments

To inhibit transcription, U2OS cells were treated with the indicated doses of Actinomycin D (Cayman Chemicals, 11421-5), DRB (Cayman Chemicals, 10010302-10) and Triptolide (Sigma-Aldrich, T3652) for the indicated time. TK6 cells were inhibited using 200μM of DRB for 3h. Efficiency of inhibition was assessed by monitoring nascent RNA levels by testing 5-Ethynyl Uridine (EU) incorporation using the Click-iT™ RNA Alexa Fluor™ 488 Imaging Kit (ThermoFisher, Scientific). EU was added to the cell media, at the final concentration of 1 mM, 1 hour before fixation. To inhibit replication cells, U2OS were treated with 20 μM Aphidicolin (Sigma-Aldrich, A4487) for 3h or TK6 cells were treated with 8 μM Aphidicolin for 3h. Efficiency of inhibition was assessed by monitoring EdU incorporation using the Click-iT™ EdU Alexa Fluor™ 488 Imaging Kit (ThermoFisher, Scientific). EdU was added to the culture media at the final concentration of 10mM, 2 hours prior to fixation.

### Immunofluorescence

For immunofluorescence, adherent cells were grown in 96 or 384 well plates (Perkin Elmer 6005550, 6057300) that were coated with poly-L-lysine (PLL, Sigma-Aldrich), were fixed in 4% formaldehyde for 10 min and washed three times with PBS. Suspension cells were gently spun down on multiwell plates and a same volume of 8% PFA was added for 20min (final concentration of 4% PFA). Cells were permeabilized with 0.3% TritonX-100 in PBS and washed twice with PBS Cells were treated with blocking buffer (3% BSA in PBS) for 1 hour before incubation with primary antibodies overnight. Cells were then washed three times in PBS containing 0.1% Tween (PBS-T) and were incubated for 1 h with fluorescently labeled secondary antibodies. After three PBS-T washes, DNA was stained with Hoechst 33342 (Sigma-Aldrich).

Primary antibodies used in this study: γH2AX (Millipore 05-636, 1:1000), Phospho RPA32(S4/S8) (Bethyl A30-245A, 1:2000), TOP2A (Santa Cruz 166934, 1:200), TOP2B (Santa Cruz 13059, 1:100).

Secondary antibodies used in this study: Alexa Fluor 568 anti-mouse IgG (Invitrogen A10037, 1:1000), Alexa Fluor 488 anti-rabbit IgG (Invitrogen A11034, 1:1000), Alexa Fluor 568 anti-rabbit IgG (Invitrogen A11011, 1:1000), Alexa Fluor 488 anti-mouse IgG (Invitrogen A-21202, 1:1000). Images were acquired by “Opera Phenix High Content Screening System” (PerkinElmer) and were analyzed by automated “Harmony High Content Imaging and Analysis Software” (PerkinElmer) as indicated below.

### siRNA knockdown

Reverse transfection of U2OS, Cal51, Hela, and MCF-7 cells with siRNAs was carried out by Lipofectamine RNAiMAX (Invitrogen 13778075) and 20nM of indicated siRNAs. 72h later transfected cell lines were challenged with Etoposide for the indicated times and doses. All siRNAs used in this study were purchased from ThermoFischer: siRNA no target (Silencer Selected negative control, #4390843); siTOP2A (Silencer Selected Validated, #4390824, siRNA ID #s14309); siTOP2B (Silencer Selected Validated, #4390824, siRNA ID #s108).

### Western blotting

Total cell lysates were either prepared by directly lysing cell pellets in SDS-page loading buffer or, for comparative analysis, prepared by lysis in RIPA buffer followed by protein concentration determination using Bradford reagent (Bio-Rad 5000205). Samples were subjected to electrophoresis on pre-cast 4%–15% gels (Bio-Rad 4561086, 4561085DC) and transferred to PVDF membranes (Merck, IPFL00010) as previously shown (Roukos et al., 2007; Stathopoulou et al., 2012). Immunodetection was performed incubating overnight at 4°C with primary antibodies diluted in PBS containing 5% milk (Sigma-Aldrich, 70166). Membranes were incubated with secondary peroxidase-coupled antibodies at room temperature for 1 hr. ECL-based chemiluminescence was detected with WesternBright Chemilumineszens Substrat Sirius (Biozym Biotech, 541021) following manufacturer’s instructions.

Primary antibodies were used at the indicated dilutions: TOP2A (G-6, Santa Cruz 166934, 1:200), TOP2A (C-15, Santa Cruz 5346, 1:200), TOP2B (H-286, Santa Cruz 13059, 1:200), TOP2B (A-12, Santa Cruz 365071, 1:200), α-Tubulin (Sigma T5168, 1:2000), MRE11 (Cell Signaling Technology 4895, 1:1000), GFP (Santa Cruz 9996, 1:1000), RAD54 (Santa Cruz, 4E3, 53433, 1:1000), LIG4 (Abcam 193353, 1:1000), TDP2 (Bethyl Laboratories A302-737A, 1:1000), CtIP (Bethyl Laboratories A300-488A, 1:1000). Secondary antibodies were used as follows: anti-mouse IgG HRP-linked (Cell Signaling Technologies 7076, 1:1000), anti-rabbit IgG HRP-linked (Cell Signaling Technologies 7074, 1:1000), anti-goat IgG HRP-linked (Santa Cruz 2354, 1:1000).

### shRNA knockdown

Lentiviral production was carried out by transfecting HEK293T cells by XtremeGene (Sigma-Aldrich, 06366236001), packaging vectors (psPax2, pMD2.G) and lentiviral vectors (shown below) in ratio 2:1:4. 16 hours later the media was substituted with complete DMEM supplemented with 30% FBS. Supernatants were collected 72 h later, filtered and concentrated 100x (Lenti-X™ Concentrator, Takara Bio 631232). Transduction of 10^6^ recipient cells was performed by spin infection with the concentrated virus and 5μg/mL polybrene (Merck TR-1003-G) at 800g for 30min. Seven days later, infected cells were challenged with Etoposide for 4h and then fixed with 4% PFA. Immunostaining and imaging were performed as before, and γH2AX levels were quantified in GFP positive cells expressing the shRNAs by using the Harmony High Content Imaging and Analysis Software (PerkinElmer). Vectors expressing shRNAs were purchased from Origene: shRNA TOP2A-1 (pGFP-C-shLenti TL308699A), shRNA TOP2A-2 (pGFP-C-shLenti TL308699C), shRNA TOP2B-1, pGFP-C-shLenti TL308698A), shRNA Top2B-2, (pGFP-C-shLenti TL308698C), shRNA scramble, (pGFP-C-shLenti shRNA Vector TR30021).

### Cell cycle synchronization and release

TK6 cells were arrested using 200μM mimosine (Sigma-Aldrich M0253) or 10μM RO-3306 (Selleckchem S7747) for 18h. After washing with PBS cells were released intro fresh medium containing the other inhibitor, respectively, and either DMSO or Etoposide (20μM or 30μM, Sigma-Aldrich E 1383) for 4h. Then, cells were washed with PBS and maintained in fresh medium in the presence of the same cell cycle inhibitor as during the etoposide treatment. After 24h, cells were spotted onto PLL coated coverslips, fixed in 4% PFA for 20 minutes and analyzed by C-Fusion 3D (see below). Cell cycle distribution was analyzed by measuring the DNA content based on Hoechst intensity as previously shown (Roukos et al., 2015) at the indicated time points.

### Fluorescence *in situ* hybridization (FISH)

Three-dimensional FISH probes were generated from bacterial artificial chromosomes (BACs from BACPAC resources center, CHORI or Thermo Scientific) by direct labeling via nick translation with fluorescently labeled dUTPs (<u>Green</u>: Chromatide AlexaFluor 488-5-dUTP, <u>Red</u>: 568-5-dUTP from Life Technologies; <u>FarRed</u>: AlexaFluor 647-AHA-dUTPfrom Thermo Fisher Scientific and <u>Blue</u>: CF405S-dUTP from Biotium) using a nick translation kit (Abbott Molecular) according to manufacturer’s instructions. For a detailed description of the BAC ID and the probed genomic locations see Table S1. The sequence specificity of all probes was verified by PCR.

For 3D FISH, cells were plated on glass poly-L-lysine coated coverslips (13 mm in diameter, 170 μm thick, Marienfeld Superior) in 24-well plates. The plates were spun at 500g in a swing out rotor centrifuge (Heraeus Multifuge X3, Thermo Fisher Scientific) for 20 seconds. After fixation in 4 % paraformaldehyde/phosphate-buffered saline (PBS) (15 minutes), cells were permeabilized (20 minutes in 0.5 % saponin (Sigma Aldrich)/0.5 % Triton X-100 (Sigma Aldrich) /PBS) and incubated in 0.1 N HCl (15 minutes) with PBS washes between steps. Cells were then washed with 2× SSC and were incubated in 50 % formamide/2× SSC buffer (30 minutes). Probe mixes (80 ng of each probe, 3 μg COT1 DNA (Roche) and 20 μg tRNA (Ambion) were ethanol precipitated and resuspended in 7 μl hybridization buffer (10 % dextran sulfate, 50 % formamide, 2× SSC, and 1 % Tween-20) and added to each coverslip. Denaturation of cells and probes was at 85 °C for 5 minutes and hybridization in a humidified chamber overnight at 37 °C. Excess probe was removed by three 5-minute washes in 1× SSC at 45 °C, followed by three 5-minute washes in 0.1× SSC at 45°C. Coverslips were mounted on glass slides (neoLab Migge GmbH) in DAPI-containing or not Vectashield mounting medium (Vector Laboratories, Burlingame, CA, USA) and sealed by picodent twinsil (Picodent 1300 1000).

### High-Throughput Imaging

Images were acquired with a spinning disk Opera Phenix High Content Screening System (PerkinElmer), equipped with four laser lines (405nm, 488nm, 568nm, 640nm). Images of FISH experiments to calculate 3D distances were acquired in confocal mode using a 40X water objective lens (NA 1.1) and two 16 bit CMOS cameras (2160 by 2160 pixels), with camera pixel binning of 2 (corresponding to 300nm pixel size). In general, for each sample 14 z-planes separated by 0.5 μm were obtained for a total number of at least 40 randomly sampled fields, which acquired in total per condition an average of 10000 cells.

Cell cycle staging of individual cells was performed by fluorescence microscopy as previously reported (Roukos et al., 2015). Briefly, for each sample 3 z-planes separated by 1 μm were obtained by using the wide-field mode of the Opera Phenix using the 20x water objective lens (NA 1.0) for a total number of at least 30 randomly sampled fields, which acquired in total per condition an average of 10000 cells.

Images of immunofluorescence samples were acquired in confocal mode of the Opera Phenix microscope using the 20x (NA 1.0) or 40x (NA 1.1) water objective lens. In general, 5 z-planes of all channels were acquired and maximum projected images were then used to measure the mean average intensity of fluorescence or number of detected foci (see next session).

### Image analysis

#### Segmentation, spot detection and intensity measurements

Image analysis was performed in the Harmony High-Content Imaging and Analysis Software (version 4.4, PerkinElmer) using standard and customized building blocks. Nuclei were segmented based on the DAPI signal or the background signal of the Red/ AlexaFluor 568 staining depending on the channel availability, using the algorithms A for suspension cell lines and C for adherent cell lines. Cells in the periphery of the image were excluded from further analysis. Mean intensity measurements were performed in maximum projected images. Custom made R scripts (<u>https://www.R-project.org/</u>) and GraphPad 7.02 were used to plot the exported upon image analysis in Harmony single cell level data. Box plots of immunofluorescence analysis show the interquartile range with whiskers representing 10-90% or 5-95% intervals (outliers are not graphically depicted but are included in statistical testing). Visualization and analysis of cell cycle profiles derived by calculating the Hoechst/DAPI nuclear staining was performed as before (Roukos et al., 2015).

#### C-Fusion 3D

FISH spots were identified in all channels in maximal projected images by using the spot detection algorithm C. Distances between all different spots of all different channels were calculated in 2D by using custom-made building blocks, while the z plane having the maximum pixel intensity of each spot was identified. Single cell information was then exported as text files and custom made scripts in R developed based on previous versions (Burman et al., 2015a), were used to calculate the Euclidean distances between different spots in 3D and to calculate frequencies of cells with chromosome breakage and/ or translocations. Correction of shifts due to chromatic aberration in z was performed by defining the pairwise shifts in z between all the different channels measured on images of a stained genomic locus by FISH using a single BAC labeled with all different color fluorophores. The value of *z* plane showing the maximum pixel intensity of each spot in every channel was corrected according the identified shift between the different channels in *z*. To determine the thresholds of physical separation and proximity, which are indicative of chromosome damage and intact loci respectively (Burman et al., 2015a; Burman et al., 2015b), we plotted the distribution of intrachromosomal distances (Green/Red and FarRed/Blue distances) in non-damaged control cells. We established a threshold of separation/proximity 4 pixels (corresponds to 1.2 μm when images were acquired with 2x binning). To eliminate cases of false or missed spot detection cases, only cells with the same number of Green and Red spots, or FarRed and Blue spots respectively, and at least 2 signals in both channels were considered in the analysis. In 4-color C-Fusion 3D, a cell with a chromosome breakage event was counted as positive, when at least one FISH Green signal that had a corresponding minimum Green/Red distance of more than the threshold (four pixels, 1.2 μm) or when a FarRed FISH signal had a corresponding minimum FarRed/Blue distance of more than the threshold. A cell was assigned positive for a chromosome translocation when three conditions were met concomitantly: 1) a Green FISH signal had a corresponding minimum Green/Red distance to the closest Red FISH signal of more than four pixels, 2) the same Green FISH signal had a minimum Green/Blue distance of more than four pixels, and 3) the minimum Green/FarRed distance of the Green FISH signal to the FarRed FISH signal was less than four pixels. Cells were assigned positive for synapsis when Green/FarRed FISH spot distances were less or equal than four pixels. In 3 color C-Fusion 3D approach, a cell positive for breakage was defined as a cell with at least one Green FISH signal that had a corresponding minimum Green/Red distance of more than four pixels. A cell with translocation was defined as a cell with at least one Green FISH signal that had a corresponding minimum Green/Red distance of more than four pixels and a concomitant Green/FarRed distance of fewer than four pixels. Euclidean distances and percentages of cells in the population showing synapsis, chromosome breakage and/ or translocations were calculated in R (https://www.R-project.org/). Plots were generated in GraphPad.

### Breaks labeling in situ and sequencing in suspension (sBLISS)

#### Cell harvesting and fixation for sBLISS

sBLISS is an adaptation of the previously published BLISS protocol (Yan et al., 2017). In sBLISS, DSB ends are labeled in cell suspensions in microcentrifuge tubes. For adherent cells, cells were washed and trypsinized following cell type-specific culturing practices. For cells grown in suspension, trypsinization was omitted and cells were washed once in PBS. Throughout the procedure, cells were pelleted with mild centrifugation (100-400g for 5 minutes, dependent on the lowest speed necessary to pellet the cells) and we made use of Protein LoBind tubes until gDNA extraction and DNA LoBind tubes for IVT and downstream library prep (Eppendorf), and low retention pipette tips. Prior to cell fixation, cell pellets were resuspended in pre-warmed PBS supplied with 10% fetal calf/bovine serum (FCS or FBS) or cell medium with the same percentage of FBS/FCS. Care was taken to ensure single-cell suspensions at this stage, either by pipetting up and down or by forcing the cells through a cell strainer. Cells were counted and diluted to reach a concentration of 10^6^ cells/ml in PBS/10%FCS. Cells were then fixed by adding 16% paraformaldehyde aqueous solution (Electron Microscopy Sciences #15710, Formaldehyde methanol-free) to a final concentration of 4% and incubating for 10 minutes on a roller shaker or tumbler at RT. Formaldehyde was quenched with 2M glycine at a final concentration of 125 mM for 5 minutes at room temperature (RT), while gently rotating, and for an additional 5 minutes on ice. Fixed cells were pelleted by centrifuging at 100-400g for 10 minutes at 4°C. After two washes with ice-cold PBS, fixed cells were kept at 4°C until BLISS template preparation.

#### BLISS template preparation

To prepare single-nucleus suspensions for tagging of DSBs by in situ ligation of BLISS adapters, 106 fixed cells were lysed for 60 minutes on ice in a lysis buffer containing 10mM Tris-HCl, 10 mM NaCl, 1 mM EDTA, and 0.2% Triton X-100 (pH 8). We then pelleted the lysed cells at RT, removed the supernatant, and permeabilized the nuclei for 60 minutes at 37°C with a second pre-warmed lysis buffer containing 10 mM Tris-HCl, 150 mM NaCl, 1 mM EDTA, and 0.3% SDS (pH 8). Then, nuclei were washed twice with pre-warmed 1x CutSmart Buffer (New England Biolabs (NEB) #B7204) supplemented with 0.1% Triton X-100 (1xCS/TX100), and DNA Double Strand Break ends (DSB ends) were blunted with NEB’s Quick Blunting Kit (NEB #E1201) according to the manufacturer’s instructions in a final volume of 100 μl for 60 minutes at RT.

Blunted nuclei were washed twice with 1x CS/TX100 before proceeding with in situ ligation of BLISS adapters (see below adapter preparation) to the blunted DSB ends. Adapter ligation was performed with 25 Weiss units T4 DNA Ligase (5 U/μl, ThermoFisher Scientific #EL0011) for 20-24h at 16°C in reaction volumes of 100 μl, according to the manufacturer’s instructions and supplemented with BSA (Thermo #AM2616) and ATP (Thermo #R0441). Per prep of 10^6^ cells, 4 μl of a 10 μM working solution of the selected BLISS adapter was ligated. Prior to use, BLISS dsDNA adapters were prepared from two complimentary HPLC-purified oligonucleotides ordered from Integrated DNA Technologies (IDT). Each dsDNA adapter contains the T7 promoter sequence for in vitro transcription (IVT), the RA5 Illumina RNA adapter sequence for downstream sequencing, an 8-nt Unique Molecular Identifier (UMI) sequence generated by random incorporation of the four dNTPs according to IDT’s ‘Machine mixing’ strategy, and an 8-nt sample barcode for multiplexing of BLISS libraries. First, sense oligos diluted to 10 μM in nuclease-free water were phosphorylated using NEB’s T4 PNK system (NEB #M0201) supplemented with ATP, after which an equimolar amount of antisense oligo was added. We used a PCR thermocycler to anneal both oligos (5 minutes at 95°C, then cooling down to 25°C in cooling steps of 1.5°C per minute) to generate a 10 μM phosphorylated dsDNA adapter.

After ligation overnight, nuclei were washed twice with 1x CS/TX100. Then we reversed crosslinks and extracted gDNA by resuspending the nuclei in 100 μl DNA extraction buffer containing 10 mM Tris-HCl, 100 mM NaCl, 50 mM EDTA, and 1% SDS (pH7.5), and supplemented with 10 μl Proteinase K (800 U/ml, NEB #P8107). Nuclei were incubated at 55°C for 14-18h while shaking at 800rpm. When clumps were still present the next morning, we added an additional 10 μl Proteinase K for 1 hour. Afterwards, Proteinase K was heat-inactivated for 10 minutes at 95°C, followed by extraction using Phenol:Chloroform:Isoamyl Alcohol 25:24:1 with 10 mM Tris, pH 8.0, 1 mM EDTA (Sigma-Aldrich/Merck #P2069) and Chloroform (Merck #1024451000), and ethanol precipitation. We sonicated the purified gDNA in 100 μl TE using a BioRuptor Plus (Diagenode) with the following settings: 30s ON, 60s OFF, HIGH intensity, 30 cycles. Sonicated samples were concentrated with Agencourt AMPure XP beads (Beckman Coulter) and fragment sizes were assessed using a BioAnalyzer 2100 (Agilent Technologies) to range from 300bp to 800bp, with a peak around 400-600bp. Sonicated and purified BLISS template was stored at −20°C until IVT and library preparation were initiated.

#### In vitro transcription (IVT) and NGS library preparation

Equal amounts (100 or 200 ng) of purified sonicated BLISS template of treated (Etoposide) and control (DMSO) samples were pooled into a single IVT reaction for linear amplification by T7-mediated transcription of the genomic ends of the BLISS-labeled DSBs. For IVT we used the MEGAscript T7 Transcription Kit (ThermoFisher #AMB13345) according to the manufacturer’s prescriptions, with the exception that transcription was carried out for 14 hours at 37°C and that Ribosafe RNAse Inhibitor (Bioline #BIO-65028) was added. Upon completion, gDNA was removed with 2 units DNase I (RNase-free) (ThermoFisher #AM2222) and amplified RNA (aRNA) was purified and concentrated using Agencourt RNAClean XP beads (Beckman Coulter). Next, oligos with the Illumina RA3 adapter sequence (Integrated DNA technologies) were ligated to the purified aRNA using T4 RNA Ligase 2, truncated (NEB #M0242) for two hours at 25°C. No RA5 adapter ligation was necessary, as this sequence was already present in each RNA molecule due to the composition of the BLISS adapter. Directly after RNA ligation, we performed reverse transcription with Reverse Transcription Primer (RTP) (Illumina sequence, ordered via IDT) and SuperScript IV Reverse Transcriptase (ThermoFisher #18090050). The manufacturer’s prescriptions were followed with the exception of incubation time, which was extended to 50 minutes at 50°C followed by heat inactivation for 10 minutes at 80°C. To prevent RNase activity, we supplemented with RNaseOUT (ThermoFisher #10777019) during RA3 ligation and reverse transcription. After completion of reverse transcription, libraries were amplified with NEBNext High-Fidelity 2x PCR Master Mix (NEB #M0541), the RP1 common primer, and a selected RPIX index primer (Illumina sequences, ordered through IDT). In total, 12 PCR cycles were completed following the manufacturer’s protocol, after which the amplified libraries were subjected to cleanup according to the two-sided AMPure XP bead purification protocol aimed to retain library sizes ranging from roughly 300-800bp. Final library profiles were assessed and quantified on a BioAnalyzer High Sensitivity DNA chip and using the Qubit dsDNA HS Assay kit (ThermoFisher #Q32851) and whenever necessary AMPure XP bead purification was repeated.

#### BLISS sequencing and data processing

Sequencing was performed in-house on a NextSeq 500 with NextSeq 500/550 High Output Kit v2 chemistry for single-end (1x76) or paired-end (2x151) sequencing, with an additional 6 cycles for index sequencing when multiple libraries were pooled. Upon completion of the run, raw sequencing reads of pooled libraries were demultiplexed based on index sequences by Illumina’s BaseSpace, after which the generated FASTQ files were downloaded. For libraries sequenced separately this step was omitted and FASTQ files were downloaded directly. As described previously (Yan et al., 2017), we applied a custom-built pipeline to keep only the reads containing the expected prefix of 8nt UMI and 8nt sample barcode using SAMtools (version 1.7) (Li et al., 2009) and scan for matches, allowing at most one mismatch in the barcode. The prefixes were clipped off and stored, and the trimmed reads per condition were aligned to the GRCh37/hg19 reference genome with BWA-MEM (version 0.7.17-r1188) (Li and Durbin, 2009). Only those reads with mapping quality scores ≥ 30 were retained. Next, we identified and removed PCR duplicates by searching for proximal reads (at most 30bp apart in the reference genome) with at most two mismatches in the UMI sequence. Finally, we generated BED files for downstream analyses, comprising a list of DSB end locations and a number of unique UMIs identified at these locations, which we refer to as ‘UMI-DSB ends’ or unique break ends.

### Bioinformatics analyses

#### Detection of BLISS hotspots

We used Macs2 (version 2.1.1.20160309) (Zhang et al., 2008) to call peaks from the BED files of UMI-DSB. First, we created one entry per UMI-DSB in the BED file, and called Macs2 assuming no background model, no cross-correlation between strands around the hotspots, and requiring Macs2 to shift the reads by −100bp and extending them 200bp in order to recreate the original fragment, with its middle position pointing to the original UMI-DSB locus. Hot spots lists from different replicates were merged using bedtools (Quinlan, 2014) (version 2.25.0; default parameters).

#### Definition of genomic regions

Gene body coordinates and transcription start sites were obtained from ENSEMBL (Release 75) (Zerbino et al., 2018) for the reference genome GRCh37 (hg19). Active genes were defined as overlapping H3K4me3 or H3K27ac histone modifications peaks or having a significant (Fisher’s exact test, P < 0.05) enrichment for RNA Polymerase II. Inactive genes were defined as the genes that did not match any of the previous criteria. Promoter regions were defined as a window of 3Kbp centered at the transcription start site of each gene. Active and inactive promoter annotations simply follow the same annotation from the gene in which it belongs to. Enhancers were defined as following: we obtained K562- and TK6-specific enhancers from the Enhancer Atlas (Gao et al., 2016) and human enhancers from FANTOM5 (Andersson et al., 2014). These two enhancer datasets were merged together, forming a comprehensive enhancer list. Active enhancers were defined as the enhancers from the comprehensive list, which overlapped H3K4me1 and H3K27ac histone modifications peaks. The remaining enhancers from the comprehensive list were defined as the inactive/ poised enhancers.

#### ENCODE data analysis

ChIP-seq (for transcription factors and histone modifications) and DNase-seq mapped reads (reference genome GRCh37; hg19, from 2016-03-06) data were downloaded from ENCODE (Davis et al., 2018) as BAM files (see Table S2). The reads from all BAM files were corrected for GC content and artificial genomic locus enrichment using BEADS (version 1.1) (Cheung et al., 2011). Furthermore, the reads from all BAM files were normalized using the reads-per-kilobase-per-million (RPKM) approach using the standard methodology (Conesa et al., 2016), when two or more datasets were being compared. When using the BAM files directly for ChIP-seq data, each read was extended downstream by 200 bp from its 5’ end to reflect the global average fragment length from the ChIP procedure (Gusmao et al., 2016). When using DNase-seq data for graphical purposes, we used the “base-overlap” signal strategy. Briefly, all reads are modified to exhibit only +/-5 bp from the 5’ end reads. For ChIP-seq data, we called peaks using Macs2 (version 2.1.1.20160309, parameters: --keep-dup auto --nomodel --nolambda --shift −100 --extsize 200 --call-summits). For DNase-seq data, all reads were first pre-processed by keeping only the 5’ end bp, which reflects the here the DNase I enzyme digested the DNA (Gusmao et al., 2016), and, then, we called peaks using Macs2 (parameters: --nomodel --nolambda --call-summits). We further filtered all peaks, for each dataset, by the Macs2-based false positive rate corresponding to their 25^th^ percentiles as recommended (Meyer and Liu, 2014).

The signal files (BIGWIG) for MNase-seq data were downloaded from ENCODE (Davis et al., 2018) and used for further analysis. sBLISS profiles in K562 cells were compared with data derived from ENCODE (K562 cells) and sBLISS analysis in the lymphoblastoid cell line TK6, was compared with ENCODE data produced from the lymphoblastoid GM12878 cells (Table S2). Data were visualized with the WashU EpiGenome Browser.

#### CTCF motives

The CTCF binding affinity motif was obtained in the Jaspar repository (Khan et al., 2018) and matched to the reference genome GRCh37 (hg19) using the package MOODS (version 1.9.3) (Korhonen et al., 2017). The motif hits were further filtered by a false positive rate of 10^−4^ using the dynamic programming estimation of the distribution of bit-scores approach as described in (Wilczynski et al., 2009). CTCF motifs with higher likehood to be bound by the CTCF protein (“active motives”), were discovered using the HINT-BC tool (version 0.11.4) (Gusmao et al., 2016) using the following approach: (1) First, DNase-seq data for K562 and TK6 cell types was obtained from ENCODE (Davis et al., 2018); (2) DNase-seq hypersensitivity sites (peaks) were called using Macs2 version 2.1.1.20160309 (parameters: --nomodel --nolambda --call-summits); (3) The final set of hypersensitivity regions was obtained by extending +-100 bp from the peak summit; (4) HINT-BC was executed using the DNase-seq data and the hypersensitivity regions as input; (5) Finally, we kept only the CTCF hits which overlapped by at least 1bp the resulting footprints from HINT-BC (see Methods section *“Overlap statistics”* for more details).

#### Chromatin Loop analyses

We obtained the chromatin intrachromosomal contacts list for K562 cells and lymphoblastoid GM12878 cells (for comparisons with TK6 cells), including both upstream and downstream loop anchors and contact-mediating CTCF binding positions, from (Rao et al., 2014). We segregated all genes into two categories: (1) Genes overlapping loop anchors – if the gene overlapped a loop anchor by at least one base pair (see Methods section *“Overlap Statistics*” for the complete details on how overlaps were performed); and (2) Genes not overlapping loop anchors – if no overlap existed between the gene and loop anchors. For the genes overlapping loop anchors, we considered a gene to be “outside a chromatin loop” if the TSS of the gene was upstream of the anchor’s CTCF (for all upstream anchors) or downstream of the anchor’s CTCF (for all downstream anchors). Similarly, all genes whose TSS were upstream of the anchor’s CTCF (for all downstream anchors) or downstream of the anchor’s CTCF (for all upstream anchors) were regarded as “inside a chromatin loop”. Given this classification the genome was segregated into four categories: (1) Genes that were in the forward and reverse strand outside of a chromatin loop; (2) genes that were in the forward and reverse strand inside of a chromatin loop; (3) genes that were exclusively in the forward strand both inside and outside a chromatin loop; and (4) genes that were exclusively in the reverse strand both inside and outside a chromatin loop. These genes were further divided into active/inactive based on the same criteria defined in the Methods section *“Definition of genomic regions*”. For clarity, we removed from these analyses all regions in which genes were overlapping in different strands. The criteria for a loop anchor to be annotated with a CTCF were either: (1) the loop anchor was already annotated with a CTCF binding site in the original list obtained in (Rao et al., 2014); or (2) the loop anchor presented a DNase-seq footprint obtained by applying the software HINT-BC (Gusmao et al., 2016) overlapping (by at least one base pair) a CTCF ChIP-seq peak obtained by applying the software Macs2 version 2.1.1.20160309 (Zhang et al., 2008) in ENCODE-derived CTCF ChIP-seq data (Davis et al., 2018). For more information, please refer to the Methods sections *“CTCF motives*” and *“Encode data analysis*”. Finally, all loop anchors which did not present a gene at least +-100Kbp from the center of their loop anchors were defined as “intergenic sites”.

#### RNA-seq data analysis

The raw data of RNA-seq primary CD34+ cells were downloaded from European Nucleotide Archive (ENA) (see Table S2). Reads were aligned on the *H. sapiens* genome assembly hg19 (GENCODE release 19) reference sequences using STAR (Dobin et al., 2013) (version 2.5.4b, parameters: --outSJfilterReads Unique --outFilterMismatchNmax 2 --outFilterMultimapNmax 10 --alignIntronMin 21 --sjdbOverhang reads_length – 1). The featureCounts program85 (version 1.5.1, parameter: -s 2) was used to count the number of reads overlapping genes. The counts were normalized by library size and gene length, then, the gene expression levels were converted to percentile-rank values.

#### GRO-seq data analysis

K562 GRO-seq data were previously generated (GSM1480325). For our purposes, we realigned the raw reads to the Illumina’s iGenomes (https://support.illumina.com/sequencing/sequencing_software/igenome.html) UCSC GRCh37 (hg19) human reference from 2016-03-06, using Bowtie (Langmead et al., 2009) (using the --best --strata –tryhard flags, allowing up to 2 mismatches and discarding reads mapping to multiple positions), after removing reads that aligned to the Human ribosomal DNA complete repeating unit (GenBank: U13369.1). We then counted the fraction of reads that fall on genes, distinguishing between exons and introns, using featureCounts from the Rsubread (Lawrence et al., 2013) package and the accompanying GTF gene annotation of the iGenomes reference genome. The counts were normalized by library size and gene length, then, the gene expression levels were converted to percentile-rank values.

#### Nascent RNA-seq (transcription factories)

Nascent RNA-seq experiment in TK6 cells was performed as previously described (Melnik et al., 2016). Raw sequencing reads were mapped to human hg19 reference genome with STAR (Dobin et al., 2013) (version 2.5.3a) and gene level counting was performed with iRNA-seq pipeline using count mode (iRNA-v1.1) (Madsen et al., 2015). The resulting read counts were normalized by reads-per-kilobase-per-million (RPKM) using the standard methodology (Conesa et al., 2016).

#### Overlap of genomic features

All overlap between two features (including genomic regions, BLISS hot spots, peaks called using MACS for transcription factors, histone modifications or open chromatin and CTCF motives) were calculated using bedtools (Quinlan, 2014) (version 2.25.0, parameters: -wa -u) -a <Regions A> -b <Regions B>) bedtools intersect -wa -u -a <Regions A> -b <Regions B> where “Regions A” are the set of intervals in which we were interested in verifying the overlap with the set of intervals “Regions B”. Overall, we defined two intervals (regions) to overlap if there is at least 1 bp overlap between them.

#### Statistical tests and corrections

To account for both linearity and non-linearity, all correlations between samples were calculated based on the Spearman’s correlation coefficient. Furthermore, since none of the datasets could be fit to a known distribution, we used the non-parametric hypothesis test Mann–Whitney–Wilcoxon in all cases of distribution comparisons. All tests were two sided and the confidence level used was 99%. The resulting p-values from all hypothesis tests were corrected for multiple alignment using the Benjamini-Hochberg method. We further corrected the reported p-values by its distribution’s false positive rate using the Longstaff and Colquhoun calculator available at <u>http://fpr-calc.ucl.ac.uk/</u> (Colquhoun, 2014). We assumed frequency equality as priors in all tests.

#### Multiple correlation test

For correlations between more than two variables, we used the nonlinear weighted least-squares method calculated with the Gauss-Newton algorithm. The summary statistics of such method provides an adjusted r^2^ statistics. The r^2^ statistics reflects the fraction of variance explained by the model; thus, it is widely used as a correlation metric. In this work, we used the square root of the adjusted r^2^ statistics as our correlation metric between three or more variables.

**Table S1.** List of BAC clones used in this study.

| **BAC name** | **Source** | **Identifier** |
| --- | --- | --- |
| MLL-5' | Thermo Fischer | CTD-2159M9 |
| MLL-3' | BACPAC Resources CHORI | RP11-59N1 |
| AF4-5' | BACPAC Resources CHORI | RP11-711J3 |
| AF4-3' | BACPAC Resources CHORI | RP11-1105H6 |
| AF6-5' | BACPAC Resources CHORI | RP11-351J23 |
| AF6-3' | BACPAC Resources CHORI | RP11-1013J5 |
| AF9-5' | BACPAC Resources CHORI | RP11-607L24 |
| AF9-3' | BACPAC Resources CHORI | RP11-459O18 |
| ENL-5' | BACPAC Resources CHORI | RP11-114A7 |
| ENL-3' | BACPAC Resources CHORI | RP11-1056A10 |
| ALK-5' | BACPAC Resources CHORI | RP11-119L19 |
| ALK-3' | BACPAC Resources CHORI | RP11-100C1 |
| NPM1-5' | BACPAC Resources CHORI | RP11-1072I20 |
| NPM1-3' | Thermo Fischer | CTD-2336J1 |

**Table S2.**
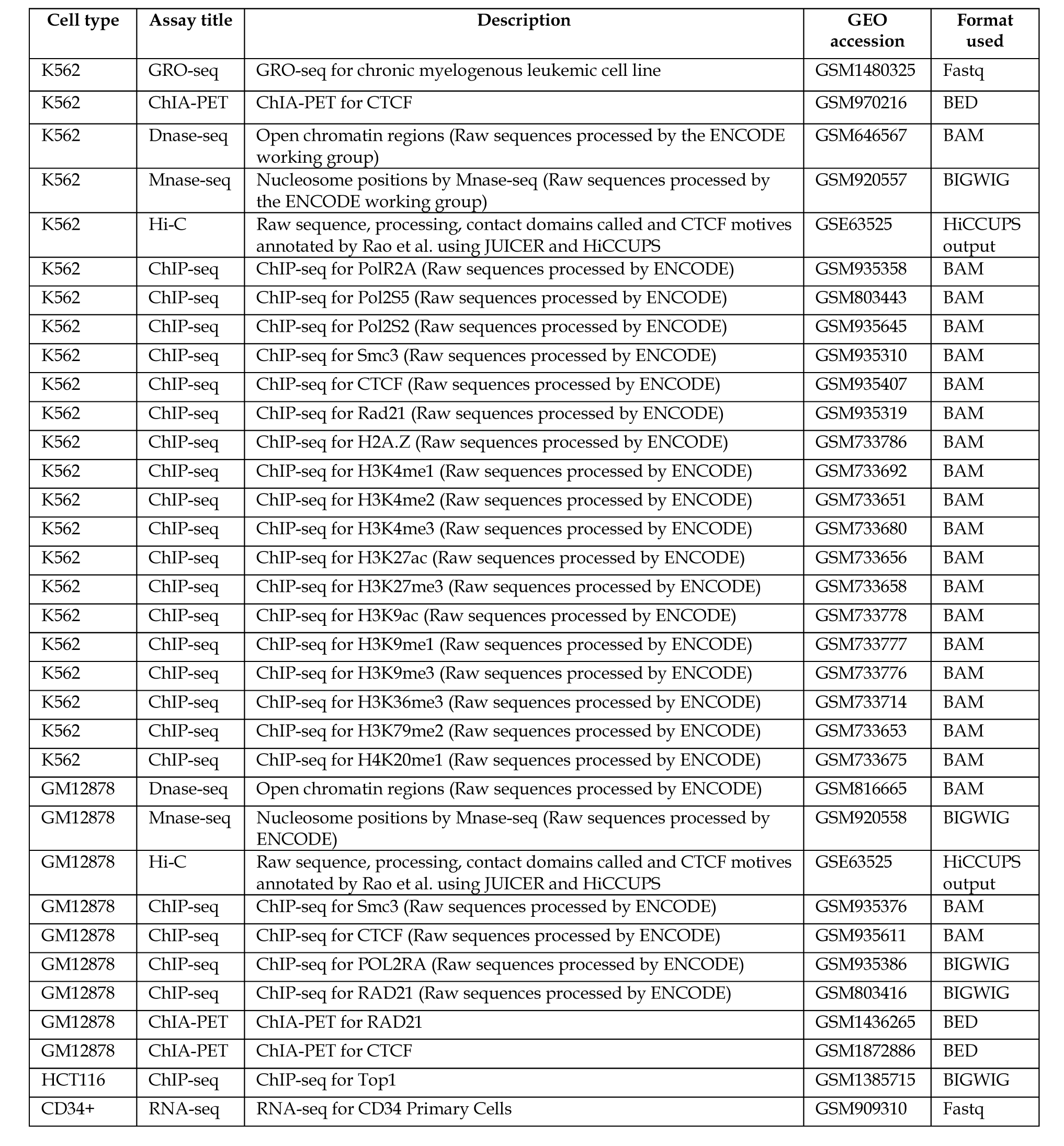
List of genomics data used in this study that have been previously generated.

## References

Allan, J.M., and Travis, L.B. (2005). Mechanisms of therapy-related carcinogenesis. Nature reviews. Cancer 5, 943–955.

Burman, B., Zhang, Z.Z., Pegoraro, G., Lieb, J.D., and Misteli, T. (2015b). Histone modifications predispose genome regions to breakage and translocation. Genes & development 29, 1393–1402.

Canela, A., Maman, Y., Jung, S., Wong, N., Callen, E., Day, A., Kieffer-Kwon, K.R., Pekowska, A., Zhang, H., Rao, S.S.P., et al. (2017). Genome Organization Drives Chromosome Fragility. Cell 170, 507–521 e518.

de Wit, E., Vos, E.S., Holwerda, S.J., Valdes-Quezada, C., Verstegen, M.J., Teunissen, H., Splinter, E., Wijchers, P.J., Krijger, P.H., and de Laat, W. (2015). CTCF Binding Polarity Determines Chromatin Looping. Molecular cell 60, 676–684.

Dobin, A., Davis, C.A., Schlesinger, F., Drenkow, J., Zaleski, C., Jha, S., Batut, P., Chaisson, M., and Gingeras, T.R. (2013). STAR: ultrafast universal RNA-seq aligner. Bioinformatics 29, 15–21.

Drynan, L.F., Pannell, R., Forster, A., Chan, N.M., Cano, F., Daser, A., and Rabbitts, T.H. (2005). Mll fusions generated by Cre-loxP-mediated de novo translocations can induce lineage reassignment in tumorigenesis. The EMBO journal 24, 3136–3146.

Gao, T., He, B., Liu, S., Zhu, H., Tan, K., and Qian, J. (2016). EnhancerAtlas: a resource for enhancer annotation and analysis in 105 human cell/ tissue types. Bioinformatics 32, 3543–3551.

Gomez-Herreros, F., Romero-Granados, R., Zeng, Z., Alvarez-Quilon, A., Quintero, C., Ju, L., Umans, L., Vermeire, L., Huylebroeck, D., Caldecott, K.W., et al. (2013). TDP2-dependent non-homologous end-joining protects against topoisomerase II-induced DNA breaks and genome instability in cells and in vivo. PLoS genetics 9, e1003226.

Gomez-Herreros, F., Zagnoli-Vieira, G., Ntai, I., Martinez-Macias, M.I., Anderson, R.M., Herrero-Ruiz, A., and Caldecott, K.W. (2017). TDP2 suppresses chromosomal translocations induced by DNA topoisomerase II during gene transcription. Nature communications 8, 233.

Goodman, L.S., Hardman, J.G., Limbird, L.E., and Gilman, A.G. (2001). Goodman & Gilman’s the pharmacological basis of therapeutics, 10th edn (New York: McGraw-Hill).

Gusmao, E.G., Allhoff, M., Zenke, M., and Costa, I.G. (2016). Analysis of computational footprinting methods for DNase sequencing experiments. Nature methods 13, 303–309.

Haffner, M.C., Aryee, M.J., Toubaji, A., Esopi, D.M., Albadine, R., Gurel, B., Isaacs, W.B., Bova, G.S., Liu, W., Xu, J., et al. (2010). Androgen-induced TOP2B-mediated double-strand breaks and prostate cancer gene rearrangements. Nature genetics 42, 668–675.

Hakim, O., Resch, W., Yamane, A., Klein, I., Kieffer-Kwon, K.R., Jankovic, M., Oliveira, T., Bothmer, A., Voss, T.C., Ansarah-Sobrinho, C., et al. (2012). DNA damage defines sites of recurrent chromosomal translocations in B lymphocytes. Nature 484, 69–74.

Heck, M.M., Hittelman, W.N., and Earnshaw, W.C. (1988). Differential expression of DNA topoisomerases I and II during the eukaryotic cell cycle. Proceedings of the National Academy of Sciences of the United States of America 85, 1086–1090.

Hoa, N.N., Akagawa, R., Yamasaki, T., Hirota, K., Sasa, K., Natsume, T., Kobayashi, J., Sakuma, T., Yamamoto, T., Komatsu, K., et al. (2015). Relative contribution of four nucleases, CtIP, Dna2, Exo1 and Mre11, to the initial step of DNA double-strand break repair by homologous recombination in both the chicken DT40 and human TK6 cell lines. Genes to cells : devoted to molecular & cellular mechanisms 20, 1059–1076.

Hoa, N.N., Shimizu, T., Zhou, Z.W., Wang, Z.Q., Deshpande, R.A., Paull, T.T., Akter, S., Tsuda, M., Furuta, R., Tsutsui, K., et al. (2016). Mre11 Is Essential for the Removal of Lethal Topoisomerase 2 Covalent Cleavage Complexes. Molecular cell 64, 580–592.

Hsiang, Y.H., Wu, H.Y., and Liu, L.F. (1988). Proliferation-dependent regulation of DNA topoisomerase II in cultured human cells. Cancer research 48, 3230–3235.

Ju, B.G., Lunyak, V.V., Perissi, V., Garcia-Bassets, I., Rose, D.W., Glass, C.K., and Rosenfeld, M.G. (2006). A topoisomerase IIbeta-mediated dsDNA break required for regulated transcription. Science (New York, N.Y) 312, 1798–1802.

Khan, A., Fornes, O., Stigliani, A., Gheorghe, M., Castro-Mondragon, J.A., van der Lee, R., Bessy, A., Cheneby, J., Kulkarni, S.R., Tan, G., et al. (2018). JASPAR 2018: update of the open-access database of transcription factor binding profiles and its web framework. Nucleic acids research 46, D260–D266.

Korhonen, J.H., Palin, K., Taipale, J., and Ukkonen, E.x (2017). Fast motif matching revisited: high-order PWMs, SNPs and indels. Bioinformatics 33, 514–521.

Kouzine, F., Gupta, A., Baranello, L., Wojtowicz, D., Ben-Aissa, K., Liu, J., Przytycka, T.M., and Levens, D. (2013). Transcription-dependent dynamic supercoiling is a short-range genomic force. Nature structural & molecular biology 20, 396–403.

Krivtsov, A.V., and Armstrong, S.A. (2007). MLL translocations, histone modifications and leukaemia stem-cell development. Nature reviews. Cancer 7, 823–833.

Langmead, B., Trapnell, C., Pop, M., and Salzberg, S.L. (2009). Ultrafast and memory-efficient alignment of short DNA sequences to the human genome. Genome biology 10, R25.

Lawrence, M., Huber, W., Pages, H., Aboyoun, P., Carlson, M., Gentleman, R., Morgan, M.T., and Carey, V.J. (2013). Software for computing and annotating genomic ranges. PLoS computational biology 9, e1003118.

Li, H., and Durbin, R. (2009). Fast and accurate short read alignment with Burrows-Wheeler transform. Bioinformatics 25, 1754–1760.

Li, H., Handsaker, B., Wysoker, A., Fennell, T., Ruan, J., Homer, N., Marth, G., Abecasis, G., Durbin, R., and Genome Project Data Processing, S. (2009). The Sequence Alignment/Map format and SAMtools. Bioinformatics 25, 2078–2079.

Lin, S., Luo, R.T., Shrestha, M., Thirman, M.J., and Mulloy, J.C. (2017). The full transforming capacity of MLL-Af4 is interlinked with lymphoid lineage commitment. Blood 130, 903–907.

Lovett, B.D., Lo Nigro, L., Rappaport, E.F., Blair, I.A., Osheroff, N., Zheng, N., Megonigal, M.D., Williams, W.R., Nowell, P.C., and Felix, C.A. (2001). Near-precise interchromosomal recombination and functional DNA topoisomerase II cleavage sites at MLL and AF-4 genomic breakpoints in treatment-related acute lymphoblastic leukemia with t(4;11) translocation. Proceedings of the National Academy of Sciences of the United States of America 98, 9802–9807.

Lyu, Y.L., Lin, C.P., Azarova, A.M., Cai, L., Wang, J.C., and Liu, L.F. (2006). Role of topoisomerase IIbeta in the expression of developmentally regulated genes. Molecular and cellular biology 26, 7929–7941.

Madabhushi, R., Gao, F., Pfenning, A.R., Pan, L., Yamakawa, S., Seo, J., Rueda, R., Phan, T.X., Yamakawa, H., Pao, P.C., et al. (2015). Activity-Induced DNA Breaks Govern the Expression of Neuronal Early-Response Genes. Cell 161, 1592–1605.

Madsen, J.G., Schmidt, S.F., Larsen, B.D., Loft, A., Nielsen, R., and Mandrup, S. (2015). iRNA-seq: computational method for genome-wide assessment of acute transcriptional regulation from total RNA-seq data. Nucleic acids research 43, e40.

Maede, Y., Shimizu, H., Fukushima, T., Kogame, T., Nakamura, T., Miki, T., Takeda, S., Pommier, Y., and Murai, J. (2014). Differential and common DNA repair pathways for topoisomerase I-and II-targeted drugs in a genetic DT40 repair cell screen panel. Molecular cancer therapeutics 13, 214–220.

Malik, M., Nitiss, K.C., Enriquez-Rios, V., and Nitiss, J.L. (2006). Roles of nonhomologous end-joining pathways in surviving topoisomerase II-mediated DNA damage. Molecular cancer therapeutics 5, 1405–1414.

Melnik, S., Caudron-Herger, M., Brant, L., Carr, I.M., Rippe, K., Cook, P.R., and Papantonis, A. (2016). Isolation of the protein and RNA content of active sites of transcription from mammalian cells. Nature protocols 11, 553–565.

Meyer, C., Burmeister, T., Groger, D., Tsaur, G., Fechina, L., Renneville, A., Sutton, R., Venn, N.C., Emerenciano, M., Pombo-de-Oliveira, M.S., et al. (2018). The MLL recombinome of acute leukemias in 2017. Leukemia : official journal of the Leukemia Society of America, Leukemia Research Fund, U.K 32, 273–284.

Meyer, C.A., and Liu, X.S. (2014). Identifying and mitigating bias in next-generation sequencing methods for chromatin biology. Nature reviews. Genetics 15, 709–721.

Natsume, T., Kiyomitsu, T., Saga, Y., and Kanemaki, M.T. (2016). Rapid Protein Depletion in Human Cells by Auxin-Inducible Degron Tagging with Short Homology Donors. Cell reports 15, 210–218.

Naughton, C., Avlonitis, N., Corless, S., Prendergast, J.G., Mati, I.K., Eijk, P.P., Cockroft, S.L., Bradley, M., Ylstra, B., and Gilbert, N. (2013). Transcription forms and remodels supercoiling domains unfolding large-scale chromatin structures. Nature structural & molecular biology 20, 387–395.

Nitiss, J.L. (2009). Targeting DNA topoisomerase II in cancer chemotherapy. Nature reviews. Cancer 9, 338–350.

Nora, E.P., Lajoie, B.R., Schulz, E.G., Giorgetti, L., Okamoto, I., Servant, N., Piolot, T., van Berkum, N.L., Meisig, J., Sedat, J., et al. (2012). Spatial partitioning of the regulatory landscape of the X-inactivation centre. Nature 485, 381–385.

Pendleton, M., Lindsey, R.H., Jr., Felix, C.A., Grimwade, D., and Osheroff, N. (2014). Topoisomerase II and leukemia. Annals of the New York Academy of Sciences 1310, 98–110.

Pombo, A., and Dillon, N. (2015). Three-dimensional genome architecture: players and mechanisms. Nature reviews. Molecular cell biology 16, 245–257.

Pommier, Y., Sun, Y., Huang, S.N., and Nitiss, J.L. (2016). Roles of eukaryotic topoisomerases in transcription, replication and genomic stability. Nature reviews. Molecular cell biology 17, 703–721.

Povirk, L.F. (2006). Biochemical mechanisms of chromosomal translocations resulting from DNA double-strand breaks. DNA repair 5, 1199–1212.

Quennet, V., Beucher, A., Barton, O., Takeda, S., and Lobrich, M. (2011). CtIP and MRN promote non-homologous end-joining of etoposide-induced DNA double-strand breaks in G1. Nucleic acids research 39, 2144–2152.

Quinlan, A.R. (2014). BEDTools: The Swiss-Army Tool for Genome Feature Analysis. Current protocols in bioinformatics 47, 11 12 11–34.

Rao, S.S., Huntley, M.H., Durand, N.C., Stamenova, E.K., Bochkov, I.D., Robinson, J.T., Sanborn, A.L., Machol, I., Omer, A.D., Lander, E.S., et al. (2014). A 3D map of the human genome at kilobase resolution reveals principles of chromatin looping. Cell 159, 1665–1680.

Roukos, V., Iliou, M.S., Nishitani, H., Gentzel, M., Wilm, M., Taraviras, S., and Lygerou, Z. (2007). Geminin cleavage during apoptosis by caspase-3 alters its binding ability to the SWI/SNF subunit Brahma. The Journal of biological chemistry 282, 9346–9357.

Roukos, V., and Mathas, S. (2015). The origins of ALK translocations. Frontiers in bioscience 7, 260–268.

Roukos, V., and Misteli, T. (2014). The biogenesis of chromosome translocations. Nature cell biology 16, 293–300.

Roukos, V., Pegoraro, G., Voss, T.C., and Misteli, T. (2015). Cell cycle staging of individual cells by fluorescence microscopy. Nature protocols 10, 334–348.

Roukos, V., Voss, T.C., Schmidt, C.K., Lee, S., Wangsa, D., and Misteli, T. (2013). Spatial dynamics of chromosome translocations in living cells. Science (New York, N.Y 341, 660–664.

Simsek, D., Brunet, E., Wong, S.Y., Katyal, S., Gao, Y., McKinnon, P.J., Lou, J., Zhang, L., Li, J., Rebar, E.J., et al. (2011). DNA ligase III promotes alternative nonhomologous end-joining during chromosomal translocation formation. PLoS genetics 7, e1002080.

Soni, A., Siemann, M., Grabos, M., Murmann, T., Pantelias, G.E., and Iliakis, G. (2014). Requirement for Parp-1 and DNA ligases 1 or 3 but not of Xrcc1 in chromosomal translocation formation by backup end joining. Nucleic acids research 42, 6380–6392.

Stathopoulou, A., Roukos, V., Petropoulou, C., Kotsantis, P., Karantzelis, N., Nishitani, H., Lygerou, Z., and Taraviras, S. (2012). Cdt1 is differentially targeted for degradation by anticancer chemotherapeutic drugs. PloS one 7, e34621.

Stingele, J., Bellelli, R., and Boulton, S.J. (2017). Mechanisms of DNA-protein crosslink repair. Nature reviews. Molecular cell biology 18, 563–573.

Tammaro, M., Barr, P., Ricci, B., and Yan, H. (2013). Replication-dependent and transcription-dependent mechanisms of DNA double-strand break induction by the topoisomerase 2-targeting drug etoposide. PloS one 8, e79202.

Toledo, L.I., Altmeyer, M., Rask, M.B., Lukas, C., Larsen, D.H., Povlsen, L.K., Bekker-Jensen, S., Mailand, N., Bartek, J., and Lukas, J. (2013). ATR prohibits replication catastrophe by preventing global exhaustion of RPA. Cell 155, 1088–1103.

Uuskula-Reimand, L., Hou, H., Samavarchi-Tehrani, P., Rudan, M.V., Liang, M., Medina-Rivera, A., Mohammed, H., Schmidt, D., Schwalie, P., Young, E.J., et al. (2016). Topoisomerase II beta interacts with cohesin and CTCF at topological domain borders. Genome biology 17, 182.

Wilczynski, B., Dojer, N., Patelak, M., and Tiuryn, J. (2009). Finding evolutionarily conserved cis-regulatory modules with a universal set of motifs. BMC bioinformatics 10, 82.

Willmore, E., de Caux, S., Sunter, N.J., Tilby, M.J., Jackson, G.H., Austin, C.A., and Durkacz, B.W. (2004). A novel DNA-dependent protein kinase inhibitor, NU7026, potentiates the cytotoxicity of topoisomerase II poisons used in the treatment of leukemia. Blood 103, 4659–4665.

Willmore, E., Frank, A.J., Padget, K., Tilby, M.J., and Austin, C.A. (1998). Etoposide targets topoisomerase IIalpha and IIbeta in leukemic cells: isoform-specific cleavable complexes visualized and quantified in situ by a novel immunofluorescence technique. Molecular pharmacology 54, 78–85.

Woessner, R.D., Mattern, M.R., Mirabelli, C.K., Johnson, R.K., and Drake, F.H. (1991). Proliferation-and cell cycle-dependent differences in expression of the 170 kilodalton and 180 kilodalton forms of topoisomerase II in NIH-3T3 cells. Cell growth & differentiation : the molecular biology journal of the American Association for Cancer Research 2, 209–214.

Wright, R.L., and Vaughan, A.T. (2014). A systematic description of MLL fusion gene formation. Critical reviews in oncology/hematology 91, 283–291.

Xiao, H., Mao, Y., Desai, S.D., Zhou, N., Ting, C.Y., Hwang, J., and Liu, L.F. (2003). The topoisomerase IIbeta circular clamp arrests transcription and signals a 26S proteasome pathway. Proceedings of the National Academy of Sciences of the United States of America 100, 3239–3244.

Yan, W.X., Mirzazadeh, R., Garnerone, S., Scott, D., Schneider, M.W., Kallas, T., Custodio, J., Wernersson, E., Li, Y., Gao, L., et al. (2017). BLISS is a versatile and quantitative method for genome-wide profiling of DNA double-strand breaks. Nature communications 8, 15058.

Yang, X., Li, W., Prescott, E.D., Burden, S.J., and Wang, J.C. (2000). DNA topoisomerase IIbeta and neural development. Science (New York, N.Y) 287, 131–134.

Yu, X., Davenport, J.W., Urtishak, K.A., Carillo, M.L., Gosai, S.J., Kolaris, C.P., Byl, J.A.W., Rappaport, E.F., Osheroff, N., Gregory, B.D., et al. (2017). Genome-wide TOP2A DNA cleavage is biased toward translocated and highly transcribed loci. Genome research 27, 1238–1249.

Zerbino, D.R., Achuthan, P., Akanni, W., Amode, M.R., Barrell, D., Bhai, J., Billis, K., Cummins, C., Gall, A., Giron, C.G., et al. (2018). Ensembl 2018. Nucleic acids research 46, D754–D761.

Zhang, A., Lyu, Y.L., Lin, C.P., Zhou, N., Azarova, A.M., Wood, L.M., and Liu, L.F. (2006). A protease pathway for the repair of topoisomerase II-DNA covalent complexes. The Journal of biological chemistry 281, 35997–36003.

Zhang, Y., Liu, T., Meyer, C.A., Eeckhoute, J., Johnson, D.S., Bernstein, B.E., Nusbaum, C., Myers, R.M., Brown, M., Li, W., et al. (2008). Model-based analysis of ChIP-Seq (MACS). Genome biology 9, R137.

Zhang, Y., McCord, R.P., Ho, Y.J., Lajoie, B.R., Hildebrand, D.G., Simon, A.C., Becker, M.S., Alt, F.W., and Dekker, J. (2012). Spatial organization of the mouse genome and its role in recurrent chromosomal translocations. Cell 148, 908–921.

Zhang, Y., and Rowley, J.D. (2006). Chromatin structural elements and chromosomal translocations in leukemia. DNA repair 5, 1282–1297.

Zhao, Y., Thomas, H.D., Batey, M.A., Cowell, I.G., Richardson, C.J., Griffin, R.J., Calvert, A.H., Newell, D.R., Smith, G.C., and Curtin, N.J. (2006). Preclinical evaluation of a potent novel DNA-dependent protein kinase inhibitor NU7441. Cancer research 66, 5354–5362.

